# Sex-Specific Characterization of a Novel Osteoarthritis-Induced Heart Failure Model in Mice

**DOI:** 10.1101/2025.10.16.682846

**Authors:** Pranav Prasoon, Kelly Tammen, Aravind Meyyappan, Manushri Dalvi, Sreejita Arnab, Melanie Eschborn, Roman Fischer, Matthew Kay, David Mendelowitz, John R. Bethea

## Abstract

Chronic low-grade inflammation is increasingly recognized as a key driver of heart failure (HF) progression; however, the direct contribution of systemic inflammatory disorders such as osteoarthritis (OA) remains unclear. Here, we establish a murine model of OA-induced HF using destabilization of the medial meniscus (DMM) to induce systemic inflammation and sex-specific cardiac remodeling. Longitudinal echocardiography revealed that females develop diastolic dysfunction with preserved ejection fraction, resembling HFpEF, whereas males exhibit progressive systolic impairment, consistent with a transitional HFmrEF-to-HFrEF phenotype. Morphometric and histological analyses confirmed concentric hypertrophy in females and eccentric remodeling in males. Transcriptomic profiling identified distinct molecular programs—females upregulated extracellular matrix, cytoskeletal, and calcium-handling genes, while males showed enrichment of inflammatory and immune signaling pathways. Immunoblot analyses further validated these sex-specific molecular signatures: females displayed increased ANP, BNP, Sirt1, and AMPK expression, consistent with metabolic resilience and fibrotic remodeling, whereas males exhibited elevated p38 MAPK, NF-κB, LC3B, and cleaved caspase-3, reflecting heightened inflammation, autophagy, and apoptosis. Both sexes demonstrated downregulation of mitochondrial and lipid metabolic proteins, indicating convergent energetic stress. Collectively, these findings identify OA as a systemic inflammatory driver of heart failure, delineate the molecular and proteomic basis of sex-dependent cardiac remodeling, and introduce a translational preclinical model that recapitulates the clinical heterogeneity of HFpEF and HFmrEF/HFrEF, providing a foundation for mechanistic and therapeutic exploration.

## Introduction

Heart failure (HF) remains a leading cause of morbidity and mortality worldwide, affecting over 64 million individuals globally(1). Clinically, HF is classified by left ventricular ejection fraction (LVEF) into heart failure with reduced EF (HFrEF; ≤40%), mid-range EF (HFmrEF; 41–49%), and preserved EF (HFpEF; ≥50%) (2). Epidemiologic data show that HFpEF accounts for ∼50% of cases, HFrEF ∼40%, and HFmrEF 10–20% (2). HFmrEF has emerged as a distinct and dynamic phenotype, often representing a transitional state of systolic dysfunction in which patients may regress toward HFrEF or improve toward HFpEF depending on treatment (3). While patients with HFrEF benefit from evidence-based, mortality-reducing therapies— including angiotensin receptor–neprilysin inhibitors, ACEi/ARB, β-blockers, mineralocorticoid receptor antagonists, and sodium–glucose cotransporter 2 inhibitors (SGLT2i)—therapeutic options for HFpEF and HFmrEF remain limited (4). Recent large-scale analyses demonstrate that SGLT2i significantly reduce cardiovascular death and HF hospitalizations across the EF >40% spectrum, with the strongest benefits in HFmrEF and attenuation in higher EF ranges (5). However, in HFpEF, empagliflozin reduces hospitalizations with only modest mortality benefit(6). This persistent therapeutic gap highlights the urgent need for translational strategies and novel preclinical models to guide the development of effective therapies.

Mounting evidence indicates that chronic inflammation is a central driver of HF pathophysiology (7). Systemic inflammatory states, including obesity, metabolic syndrome, autoimmune disorders, and aging, increase risk for HFpEF(8–10). Importantly, epidemiological and mechanistic studies consistently demonstrate sex-specific vulnerability: women disproportionately develop HFpEF, whereas men more frequently develop HFrEF(11, 12). These differences likely reflect divergent inflammatory, hormonal, and metabolic responses, though the precise mechanisms remain incompletely understood.

Osteoarthritis (OA) is the most common chronic joint disease, affecting more than 500 million people worldwide. Beyond its musculoskeletal burden, OA confers substantial cardiovascular risk. Large cohort studies reveal that individuals with OA have up to a three-fold higher risk of developing HF compared to the general population(13–15). This association is mediated through shared pathophysiological processes, including persistent low-grade inflammation, endothelial dysfunction, and systemic metabolic alterations (16). Despite the strong epidemiological link, the causal contribution of OA-driven inflammation to HF progression has remained poorly defined, in part due to the lack of appropriate preclinical models.

Most available HF models rely on genetic manipulation, pressure overload (e.g., transverse aortic constriction), or ischemic injury (17). While these approaches have provided critical insights into cardiac remodeling and dysfunction, they do not adequately capture the chronic inflammatory milieu observed in many human patients with HFpEF. Notably, clinical populations increasingly present with HFpEF driven by comorbid inflammatory diseases rather than isolated genetic or hemodynamic insults (18, 19). Thus, there is a pressing need for preclinical models that replicate this inflammatory phenotype and its sex-specific outcomes.

To address this gap, we developed and characterized a novel murine model in which OA serves as the initiating factor for HF. Surgical destabilization of the medial meniscus (DMM) induces OA and systemic inflammation, leading to distinct cardiac phenotypes between sexes. Female mice develop diastolic dysfunction with preserved ejection fraction, closely mirroring HFpEF, whereas male mice develop progressive systolic dysfunction consistent with HFmrEF. This divergence highlights the translational relevance of OA-induced HF as a model system, capturing the heterogeneity, sex-specificity, and inflammatory underpinnings of human HF.

In summary, OA-induced HF marks a significant advancement over traditional genetic models by directly linking a common inflammatory comorbidity to cardiac dysfunction. By employing functional measurements, histology, transcriptomics, and molecular analysis, we delved deeper into the mechanisms driving OA-induced heart failure. These analyses uncovered distinct patterns of dysfunction between males and females, underscoring the sex-specific nature of cardiac responses to chronic inflammation. This model provides a valuable platform for investigating the mechanisms of sexually specific disease progression and for assessing novel therapeutic strategies targeting inflammation-driven heart failure.

## Materials and Methods

### Ethical Statement

The animal experiments were approved by the Institutional Animal Care and Use Committee (IACUC) at George Washington University (GWU), Washington, DC, USA. They were conducted in accordance with the animal use protocol (GWU IACUC no. A2024-091) and the guidelines of the National Institutes of Health. All ethical guidelines were adhered to during the execution of this study.

### Animals

Eleven-week-old male and female C57BL/6J (Cat. 000664) mice were obtained from The Jackson Laboratory. Before the experiments, the mice were given a one-week acclimation period. They were kept in an environment with a temperature range of 23–25 °C, under a 12-hour light cycle starting at 6:00 a.m. and a 12-hour dark cycle commencing at 6:00 p.m. Humidity levels were maintained between 30–70%, and water was available ad libitum. At the study’s conclusion, the mice were humanely euthanized using a ketamine and xylazine cocktail. Tissues were immediately collected and stored appropriately for further analysis.

### Osteoarthritis-induced Heart Failure model

We performed the Surgical Induction of Osteoarthritis via DMM surgery. Osteoarthritis was induced in 12-week-old C57BL/6J male and female mice using the Destabilization of the Medial Meniscus (DMM) model, as previously described (Glasson, Blanchet et al. 2007). Mice were anesthetized with isoflurane (1–1.5 % in oxygen), and the right knee joint was shaved and sterilized with betadine and ethanol. A medial parapatellar incision was made to expose the joint capsule, and the medial meniscotibial ligament (MMTL) was transected under a surgical microscope to destabilize the medial meniscus. The capsule and skin were sutured using 6-0 absorbable sutures and surgical glue, respectively. Sham surgeries were performed on contralateral knees or control animals by exposing the joint without ligament transection. Post-operative analgesia (e.g., buprenorphine, 0.05 mg/kg, subcutaneously) was administered for 48 hours. Mice were monitored daily for signs of distress and allowed unrestricted activity until sacrifice. We performed von Frey testing to evaluate pain hypersensitivity. Pain onset in both male and female subjects was observed approximately four weeks post-DMM surgery, which subsequently resulted in osteoarthritis-induced heart failure in both sexes in later time points.

### Von Frey Testing

Von Frey testing was employed to assess mechanical hypersensitivity, serving as an indicator of a pain-like phenotype in mice (Chaplan, Bach et al. 1994). Mice were individually placed in plexiglass chambers elevated on a mesh-wire platform and were permitted to acclimate for a duration of 45 to 60 minutes prior to testing. Using the up-down method, the plantar surface of the hind paw, especially near the toe, was stimulated with von Frey filaments (Touch-test sensory evaluator) of varying diameters (force range: .02g-2g). In each trial, an intermediate filament weighing 0.16 g was applied perpendicularly to the skin, causing a slight bend for a duration of 3 seconds. A positive response, marked by the rapid withdrawal or licking of the paw within 3 seconds after the filament’s removal, while ignoring normal movements like ambulation or rearing, prompted the use of the next smaller filament. If no response was observed, the next larger filament was tested. Trials continued until four measurements were recorded following the initial change in response, whether from no response to a response or vice versa. The 50% mechanical withdrawal threshold was calculated using the statistical method outlined by Dixon (1965). For the assessment of Osteoarthritis-induced pain, the right hind paws (ipsilateral side) were utilized. The baseline withdrawal threshold of the mice was assessed one day prior to DMM surgery, and von Frey testing was conducted from 3 to 12 weeks post-DMM surgery.

### Echocardiography (ECHO)

ECHO measurements were acquired using our VisualSonics Vevo 3100, which simultaneously acquires body temperature, respiratory rate, and the ECG. A 2D parasternal long-axis view was used to measure LV outflow tract diameter. The M-mode parasternal short-axis view, at the plane of the papillary muscles, provided LV diastolic posterior wall diameter (LVPWd) and interventricular septal diameter (IVSd), LV diameter in diastole (LVDd) and systole (LVDs), LV systolic posterior-wall (LVPWs) thickness, and LV ejection fraction (EF%). These measurements will be used to calculate fractional shortening (FS). The apical four-chamber view wereused to measure velocity through the mitral valves for assessment of diastolic function, which includes early diastolic mitral inflow (E-wave), late diastolic mitral inflow (A-wave), early-to-late diastolic mitral inflow ratio (E/A ratio), and early diastolic velocity (e’). Continuous-wave Doppler of the ascending aorta were used to measure velocity and pressure through the ascending aorta, allowing the assessment of HR and cardiac output.

### Histology

#### Knee

Knee joints were drop-fixed in 10% paraformaldehyde for 72 h. Samples were decalcified with 14% EDTA for 2 weeks. Tissues were then transferred to 30% sucrose solution for 3 days then embedded in O.C.T. O.C.T embedded tissues were sectioned at 10 μm sections and stained with safranin O-fast green.

Cartilage degradation was assessed in knee joints through proteoglycan loss and fibrillation of knee tissue. All stained sections were examined using a Leica DMR microscope with a 10x objective, equipped with a Leica DFC 300 FX digital camera. Histological scoring was conducted on four to six sections per animal.

#### Heart

Heart samples were drop-fixed in 10% paraformaldehyde for 72 h. Tissues were then transferred to 30% sucrose solution for 3 days then embedded in O.C.T. O.C.T embedded tissues were sectioned at 15 μm sections and stained with Hematoxylin and Eosin (H&E). Protocol was obtained and adapted to heart tissue (20). All stained sections were examined using a Leica DMR microscope with a 10x objective, equipped with a Leica DFC 300 FX digital camera. Histological scoring was conducted on 3 sections per animal.

### Histological Scoring: (Knee and Heart)

A semiquantitative scoring system for murine histopathology, the OARSI score (21, 22) was applied and adapted to our experimental conditions. All four quadrants of the knee joint were evaluated: medial femoral condyle (MFC), lateral femoral condyle (LFC), medial tibial plateau (MTP), and lateral tibial plateau (LTP). A score from 0 to 6 was given to each quadrant of 3 serial sections per animal, for a total of 12 values per animal. The final histological scores were expressed as the average of all the individual values, and the average summed score for each experimental group was calculated. Two observers scored all the histological changes and were blinded to the specimen samples. A third observer was involved if scores differed greater by one point on the OARSI scale.

Heart: Left ventricular wall thickness was evaluated through ImageJ software. Three measurements were performed of varying regions of the left ventricular wall. This was averaged and repeated in three animals per experimental group. Chamber cavity was obtained through measuring the maximum distance between the intraventricular wall and left ventricular wall. This was repeated across three animals per experimental group.

### Western Blots

The left ventricular heart tissue was lysed in RIPA buffer (10 mM Tris-HCl pH 7.4, 1 mM EDTA, 0.5 mM EGTA, 1% NP-40, 0.1% sodium deoxycholate, 0.1% SDS, 140 mM NaCl) with added protease (Santa Cruz) and phosphatase (Biovision) inhibitors. The homogenates were kept on a rocker for 60 minutes at 4 °C, then centrifuged at ≥14,000 g at 4 °C for 15 minutes, and the resulting supernatant was collected. Protein concentrations were measured using the DC protein assay (Bio-Rad). Protein extracts were separated by sodium dodecyl sulfate polyacrylamide gel electrophoresis on 12% gels for ANP and Gal4, and 10% gels for Col1a1 and Col3a1. The gels were transferred to nitrocellulose membranes (Turbo blot,

Bio-Rad) and blocked for 2 hours in 5% bovine serum albumin (BSA) in 1x TBS-T (10 mM Tris-HCl, pH 7.5, 150 mM NaCl, 0.1% Tween-20). Nitrocellulose membranes were incubated with primary antibodies diluted in blocking solution at 4 °C overnight. The primary antibodies employed in the immunoblotting process are detailed in Supplementary Table 2. After incubation with primary antibodies, membranes were treated with horseradish peroxidase-conjugated species-specific secondary antibodies. Protein bands were detected using a chemiluminescent substrate (West Pico, ThermoFisher Scientific), imaged with ChemiDoc (BioRad), and band intensities were quantified using Image Lab software (BioRad). The total amount of protein loaded was used to normalize data through visualization with Ponceau S solution (Sigma).

### RNAseq

Total RNA was extracted from the left ventricle tissue of hearts dissected from both male and female mice (n = 3 biological replicates per group) under three experimental conditions: naïve and OA group. The extraction was performed using the RNAeasy Plus Universal Mini Kit from Qiagen [Lot no. 181018346], following the manufacturer’s protocol, which included on-column DNase digestion to remove genomic DNA contamination. RNA quality and integrity were evaluated using [Bioanalyzer 2100 / Tape Station; Agilent Technologies], and samples with an RNA integrity number (RIN) ≥ 7.0 were selected for library preparation. RNA sequencing libraries were prepared using the Illumina kit with [poly(A) selection / rRNA depletion] as per the manufacturer’s instructions. Sequencing was conducted on NovaSeq 6000 to produce [paired-end / single-end] reads of [read length; e.g., 150 bp], with an average depth of [20 M paired reads sample (40M total] reads per sample. Reads underwent quality checks using FastQC and were trimmed with [Trimmomatic / Cutadapt] to remove adapters and low-quality bases. High-quality reads were aligned to the mouse reference genome (GRCm[version]) using [alignment software; e.g., STAR aligner], and gene-level counts were quantified using [featureCounts / HTSeq-count] based on Ensembl. Differential expression analysis was performed in R (version 4.4.1) using the DESeq2 package (version 1.44.0). Raw counts were normalized using DESeq2’s median-of-ratios method, and statistical significance was determined using the Wald test with Benjamini–Hochberg adjustment for multiple testing. Genes with an adjusted p-value < 0.05 and |log2 fold change| > 1 were considered significantly differentially expressed. Data visualization included principal component analysis (PCA) plots, volcano plots, and heatmaps, generated using DESeq2 and associated R packages (e.g., ggplot2, pheatmap).

### GO functional enrichment

Differentially expressed genes (DEGs) were analyzed for Gene Ontology (GO) enrichment using the cluster Profiler R package with the org.Mm.eg.db mouse annotation database. Gene symbols were mapped to Entrez IDs with AnnotationDbi, and significant genes were defined as those with a raw p-value < 0.05. GO enrichment was performed across Biological Process (BP), Cellular Component (CC), and Molecular Function (MF) categories, and enriched terms were visualized with dot plots, where dot size represented the number of genes per term and color indicated p-value. To further assess gene–term relationships, cnet plots were generated with enrich plot in a circular layout, highlighting shared genes among the top 12 enriched categories for each ontology.

## Result

### DMM surgery induces mechanical hypersensitivity and cartilage degeneration in male and female mice

To establish the OA model, we subjected male and female C57BL/6 mice to destabilization of the medial meniscus (DMM) and evaluated mechanical sensitivity and joint pathology. Behavioral testing revealed a progressive reduction in mechanical withdrawal thresholds beginning at 3 weeks post-surgery, consistent with the onset of mechanical allodynia. Both sexes developed significant hypersensitivity in the ipsilateral hind paw relative to contralateral controls (Fig. 1B–C).

**Figure 1.**
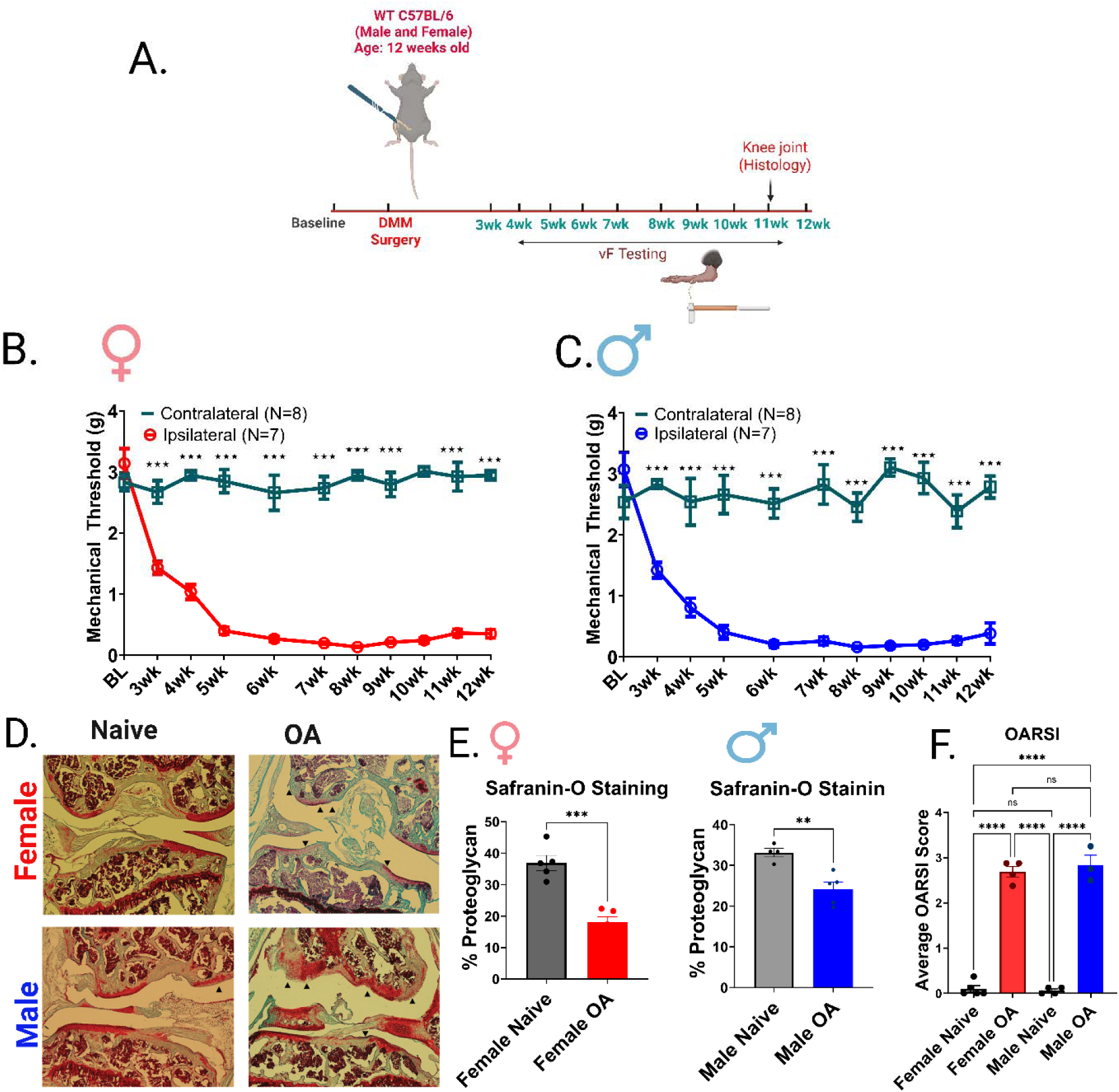
Development of mechanical hypersensitivity and cartilage degradation following DMM surgery in male and female mice. (A) Experimental timeline schematic. Twelve-week-old male and female C57BL/6 mice underwent destabilization of the medial meniscus (DMM) surgery. Mechanical thresholds were assessed weekly from 3–12 weeks post-surgery using von Frey (vF) testing. Knee joint histology was performed at 12 weeks post-surgery. (B–C) Mechanical withdrawal thresholds over time in (B) female and (C) male mice. Ipsilateral (surgery side) paws exhibited progressive and significant mechanical hypersensitivity compared to the contralateral side of the paw (control). Data are mean ± SEM; n = 7–8 per group. Two-way ANOVA with Sidak’s post-hoc test; *p < 0.05, **p < 0.01, ***p < 0.001. (D) Representative Safranin-O–stained knee joint sections from naïve and OA groups in female (top row, red label) and male (bottom row, blue label) mice. OA groups show a marked reduction in2Osteoarthritis (OA) induces sex-specific Heart Failure  proteoglycan staining compared to naïve. (E) Quantification of Safranin-O staining demonstrating significant proteoglycan loss in both female and male OA groups relative to naïve controls. Data are mean ± SEM; n = 4–5 per group. One-way ANOVA with Tukey’s post-hoc test; **p < 0.01, ***p < 0.001. (F) Quantification of cartilage degeneration using OARSI scoring. Both female and male OA groups exhibited significantly higher scores compared to naïve controls. Data are mean ± SEM; n = 4–5 per group. One-way ANOVA with Tukey’s post-hoc test; ****p < 0.0001.

Histological analyses of knee joints at 12 weeks post-surgery provided clear evidence of cartilage damage. Safranin-O staining, which labels proteoglycan content in articular cartilage, demonstrated substantial depletion in DMM-operated joints compared to naïve controls (Fig. 1D). Loss of Safranin-O staining intensity corresponded with areas of surface fibrillation and structural disruption, indicative of progressive cartilage degeneration. Quantitative analysis revealed significant reductions in safranin-O-positive regions across both sexes, confirming widespread proteoglycan loss as a hallmark of OA pathology (Fig. 1E). Consistent with these findings, OARSI scoring further substantiated cartilage degeneration in female and male mice. Compared to naïve controls, DMM-operated joints exhibited robustly elevated OARSI scores, reflecting the combined severity of proteoglycan depletion, cartilage surface irregularity, and structural breakdown (Fig. 1F). Importantly, this scoring method provided a semiquantitative assessment that paralleled the histological observations, validating the reproducibility and severity of the OA phenotype induced by DMM surgery in mice.

Together, these results demonstrate that DMM surgery induces pain hypersensitivity and significant structural joint pathology in both male and female mice. The concordance between the behavioral, histological, and quantitative scoring metrics underscores the reliability of the DMM model for studying OA-associated pain and degeneration in both sexes. Notably, the convergence of mechanical allodynia, proteoglycan depletion, and high OARSI scores indicates a local inflammatory environment triggered by OA, which is a key driver of downstream cardiovascular dysfunction explored in subsequent figures.

### Develop a mouse model that mimics human Heart Failure

Heart failure (HF) is a multifactorial disease in which inflammation is a central mediator of disease progression, and women are disproportionately affected. To model the clinical spectrum of HF associated with osteoarthritis (OA), we established a murine model in which OA-driven inflammation promotes sex-specific cardiac remodeling. Echocardiographic and morphometric assessments revealed that OA induction precipitated distinct HF phenotypes in females and males, paralleling human HFpEF and HFrEF, respectively.

### Female mice develop HFpEF

Representative echocardiographic tracings from female OA mice demonstrated preserved systolic function but impaired diastolic relaxation compared with that in naïve controls. Quantitative analysis confirmed that the ejection fraction remained stable at both 8- and 16-weeks post-OA (Fig. 2C). However, diastolic indices, E/A and E/e’ ratios, revealed progressive dysfunction, with the onset of significant changes, a decrease, and an increase in the ratios, respectively, at 8 weeks (Fig. 2C). At 16 weeks, both the E/A and E/e’ ratios were significantly elevated, indicating progression into the later stages of HFpEF (Fig. 2C). Prolonged isovolumetric relaxation time (IVRT) (Fig. 2I) further established the impaired ventricular relaxation. In addition, morphometric analysis at 16 weeks revealed an increased heart weight-to-tibia length ratio (HW/TL³) (Fig. 2 K), which was consistent with concentric hypertrophy. Together, these findings establish a phenotype of HFpEF in females, characterized by preserved ejection fraction, diastolic dysfunction, and concentric remodeling of the left ventricle.

**Figure 2.**
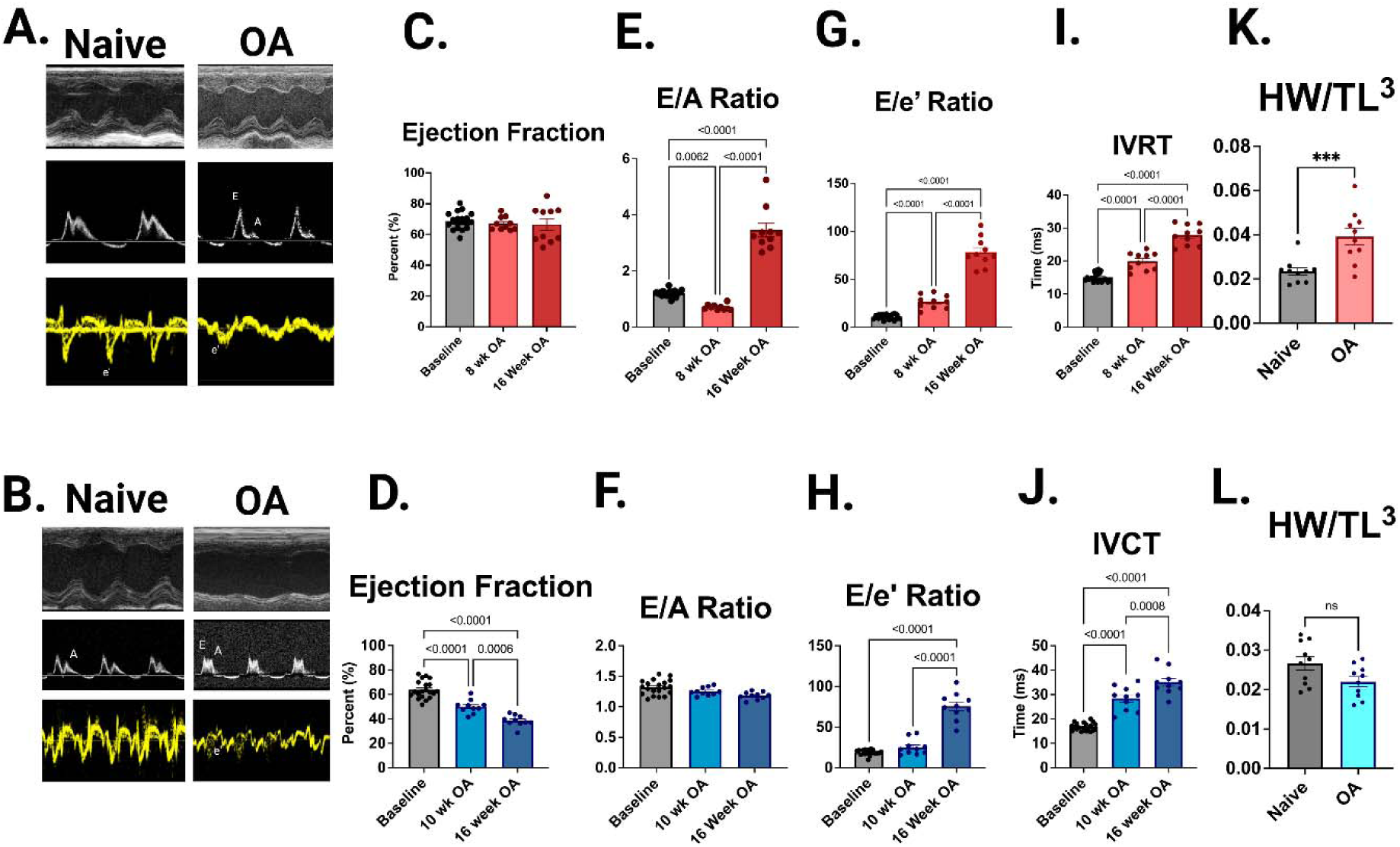
Osteoarthritis (OA) induces sex-specific Heart Failure phenotypes in mice. **A–B**, **Echocardiographic tracings**. Representative M-mode, pulse-wave Doppler (E/A ratio), and tissue Doppler (E/e′) recordings from naïve and OA mice illustrate sex-specific alterations in cardiac performance. Female OA mice (A) displayed preserved systolic function with impaired diastolic relaxation, consistent with heart failure with preserved ejection fraction (HFpEF). Male OA mice (B)This aligns with systolic dysfunction and a phenotype resembling heart failure with mid-range ejection fraction (HFmrEF), which represents a transitional phase of heart failure with reduced ejection fraction (HFrEF). The onset of heart failure of preserved ejection fraction (HFpEF) was observed at 8 weeks in females and heart failure of reduced ejection fraction (HFrEF) at 10 weeks in males. C–I, Female cardiac function (HFpEF). Quantitative analyses showed that females-maintained ejection fraction at 8- and 16-weeks post-OA compared to baseline (C), but developed progressive diastolic dysfunction characterized by elevated E/A (E) and E/e′ ratios (G). Relaxation abnormalities were evident with significantly prolonged isovolumetric relaxation time (IVRT) (I). Together, these features establish diastolic impairment with preserved systolic function, consistent with HFpEF. **D–J**, **Male cardiac function (HFmrEF)**. Males demonstrated a significant decline in ejection fraction beginning at 10 weeks and further reduction by 16 weeks post-OA (D). Diastolic indices, including elevated E/A (F) and E/e′ ratios (H), indicated impaired filling pressures. In addition, isovolumetric contraction time (IVCT) was significantly prolonged (J), reflecting impaired contractility. These changes indicate progressive systolic dysfunction characteristic of HFrEF. K–L, Morphometric assessment of hypertrophy. Female OA mice exhibited a significant increase in heart weight-to-tibia length ratio (HW/TL³) (K), consistent with concentric hypertrophy observed in HFpEF. In contrast, males showed no significant change in HW/TL³ (L), aligning with eccentric remodeling and wall thinning typically seen in HFrEF. **Overall interpretation**. OA drives sex-specific cardiac remodeling: females develop HFpEF with preserved systolic function but impaired diastolic relaxation and concentric hypertrophy, whereas males progress to HFmrEF with systolic decline and eccentric remodeling. Data are presented as mean ± S.E.M.; statistical analysis by one-way ANOVA with post hoc multiple comparisons. *P < 0.05, **P < 0.01, ***P < 0.001. Group sizes: n=8–10 for echocardiographic measures; n=5 for morphometry.

### Male mice develop HFmrEF

In contrast, male mice with OA exhibited a progressive decline in systolic performance. Representative echocardiography showed reduced contractility relative to both naïve males and baseline recordings, and quantification revealed a significant drop in ejection fraction beginning at 10 weeks and further decline at 16 weeks post-OA (Fig. 2D). Males did not show changes in the E/A ratio at 10 and 16 weeks but did show an increase in E/e’ at 16 weeks (Fig. 2F, H). Importantly, the isovolumetric contraction time (IVCT) was significantly prolonged (Fig. 2 J), reflecting the impaired contractile dynamics. Unlike females, morphometric assessment showed no increase in HW/TL³ (Fig. 2 L), consistent with the eccentric remodeling and wall thinning typical of HFmrEF.

These results demonstrate that OA induces sex-specific cardiac phenotypes in mice. Female mice develop HFpEF with preserved systolic function, impaired diastolic relaxation, and concentric hypertrophy. In male mice, the development of systolic dysfunction, impaired contraction, and eccentric remodeling leads to a phenotype that closely resembles heart failure with mid-range ejection fraction (HFmrEF). This condition serves as a transitional stage towards heart failure with reduced ejection fraction (HFrEF), marked by a decline in systolic function. This OA-driven model recapitulates the clinical diversity of HF syndromes, establishing it as a relevant inflammation-driven system for studying sex-specific mechanisms of heart failure progression.

### Sex-specific cardiac remodeling in OA mice revealed by histological analysis

Hematoxylin and eosin (H&E) staining of left ventricular (LV) sections revealed sex-specific remodeling patterns after OA induction. In female rats, OA hearts exhibited concentric hypertrophy, with reduced LV chamber diameter and increased wall thickness, consistent with a heart failure with preserved ejection fraction (HFpEF)–like phenotype (Fig. 3A–B). In contrast, males displayed eccentric hypertrophy, characterized by enlarged LV chambers and thinner ventricular walls, indicative of a heart failure with reduced ejection fraction (HFmrEF)-like phenotype (Fig. 3C–D). Quantification of the chamber diameter and wall thickness (Fig. The results shown in Figure 3E and F) confirm these divergent remodeling responses, with significant changes observed in females but not in males, underscoring sex-specific susceptibility to OA-induced cardiac remodeling.

**Figure 3.**
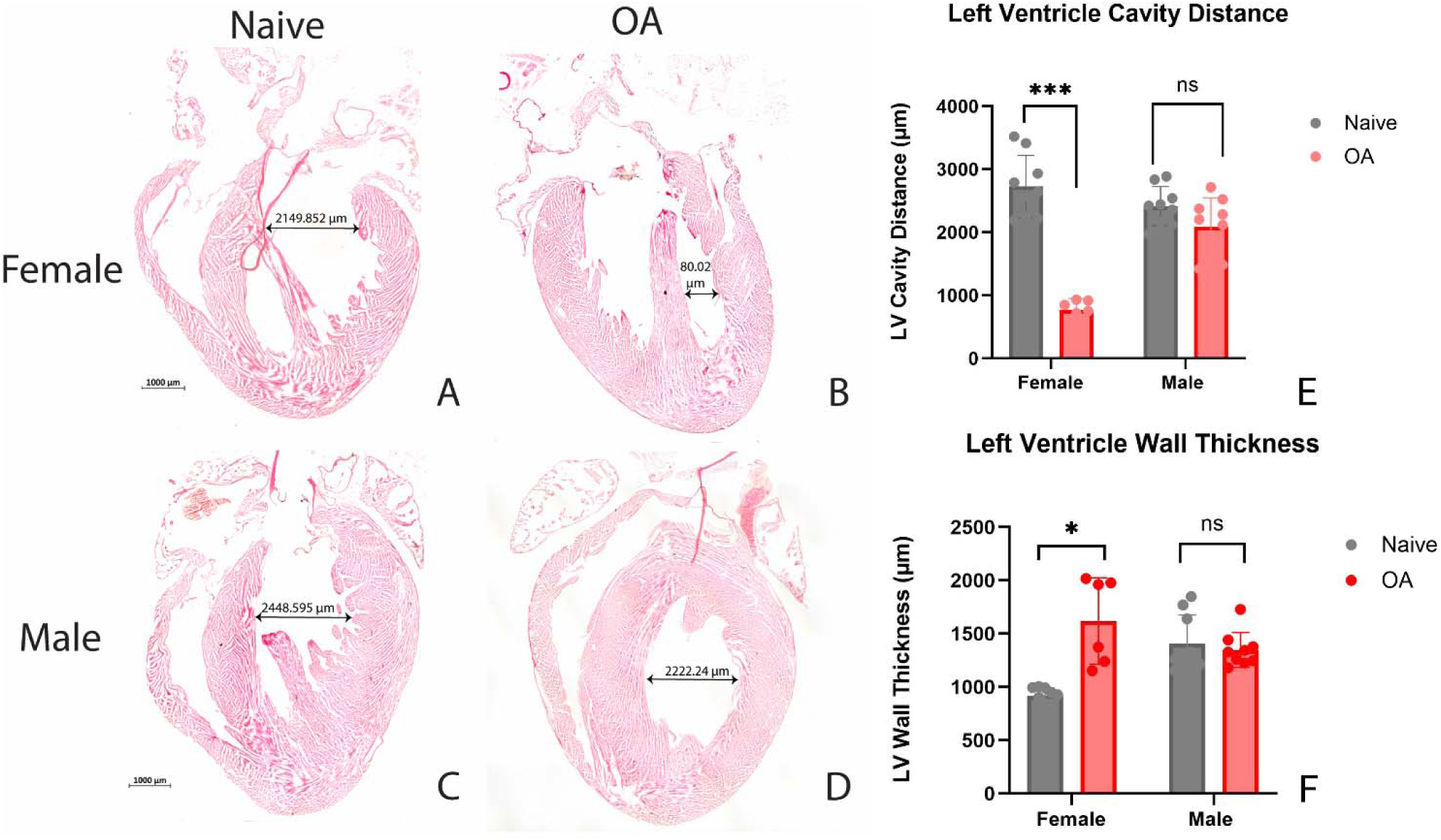
Histological assessment of left ventricular remodeling in OA mice. Representative hematoxylin and eosin (H&E)–stained heart sections from naïve and OA mice demonstrate sex-specific patterns of cardiac hypertrophy. (A, B) Naïve and OA female hearts, respectively. (C, D) Naïve and OA male hearts, respectively. (B) Female OA hearts exhibit concentric hypertrophy, characterized by reduced left ventricular (LV) chamber diameter and increased wall thickness, consistent with diastolic dysfunction and a heart failure with preserved ejection fraction (HFpEF)–like phenotype. In contrast, (D) male OA hearts exhibit eccentric hypertrophy; however, we did not observe changes in LV chamber diameter or ventricular wall thickness, despite the presence of an enlarged LV chamber diameter and thinner ventricular walls. This aligns with systolic dysfunction and a phenotype resembling heart failure with mid-range ejection fraction (HFmrEF), which represents a transitional phase of heart failure with reduced ejection fraction (HFrEF). (E, F) Quantification of LV chamber diameter and wall thickness from histological sections, displayed as bar diagrams, further highlights the sex-specific patterns of concentric versus eccentric hypertrophy. Scale bars = 1000 μm

### Transcriptomic Profiling in OA-Induced Heart Failure

RNA sequencing of left ventricular tissue revealed robust transcriptional reorganization with clear sex-specific differences in gene expression. Heatmap clustering (Fig. 4A–C) highlighted the distinct gene regulation of cytoskeletal, immune, and inflammatory pathways between the groups. Volcano plots (Fig. 4D–F) further underscored the sex divergence. **Cytoskeletal genes (Fig. 4D), sarcomeric genes such as TTN, Neb, and MYH7B were strongly upregulated in females**, indicating altered contractile stiffness. Notably, TTN expression increased by ∼40-fold and Neb by ∼15-fold in OA females compared with naïve controls, changes not observed in males. In contrast, male hearts showed upregulation of cytoskeleton-related genes involved in contractility and transport (KIF6 and AMPH), suggesting changes in vesicle and ion movement related to cardiac contractility. **Immune gene (Fig. 4E):** Females displayed induction of immune effectors, including the upregulation of Galectin-4 and Fcα/μ receptor, and the downregulation of CD74/CXCR4. Female mice also showed a large increase in immunoglobulin transcripts, suggesting increased B cell activity. In contrast, males showed enrichment of immunoregulatory transcripts (CD96, LIF, IL12α, and Apoa1) and chemokine receptors (CXCR2 and CXCR5), reflecting divergent immune trajectories. **Inflammatory gene (Fig. 4F):** Female hearts exhibited broad induction of inflammatory mediators, including transcription factors (Cebpb and Nfkb1), macrophage/monocyte markers (Ly6g6c and Cd68), cytokine receptor Il10Ra, and chemokines (Ccl2 and Cxcl10). In males, induction was more selective, with elevated Alox15, Cxcr2, Cxcr5, Il12a, complement components (Cfd, C4b), and inhibitory regulators (Tnfaip3, Nfkbiz), consistent with an immunoregulatory bias in males.

**Figure 4:**
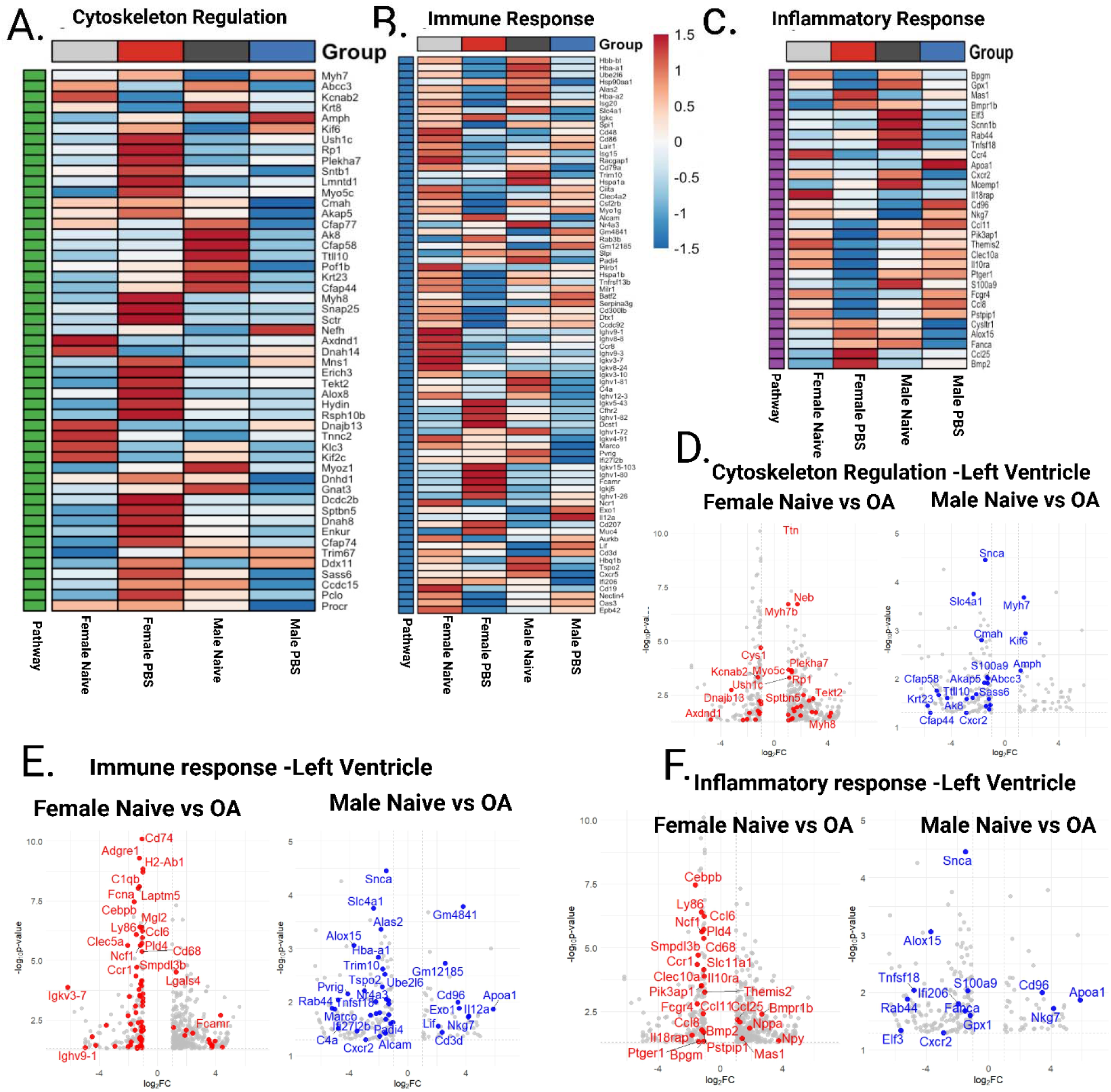
Transcriptomic analysis of left ventricular tissue reveals distinct cytoskeletal, immune, and inflammatory profiles in males and females following novel OA-induced heart failure model. (A–C) Heatmaps illustrate differential gene expression in left ventricular (LV) tissue, highlighting pathways related to cytoskeletal remodeling, immune regulation, and inflammatory signaling. Samples were clustered based on normalized expression value obtained from bulk RNA sequencing, uncovering distinct molecular profiles between heart failure phenotypes and controls. (D–F) Volcano plots display significantly dysregulated and upregulated genes in males and females, with emphasis on genes involved in fibrotic remodeling, immune activation, and pro-inflammatory signaling.

Similarly, as shown in Supplementary Figure 1, transcriptomic profiling of left ventricular tissue uncovered widespread remodeling of genes involved in lipid metabolism and mitochondrial function. Heatmaps demonstrated clear clustering of experimental groups, with coordinated induction of lipid remodeling genes such as Apoa1, Elovl4, Cyp2c23, Bmpr1b, and Fut2 in OA-induced heart failure, while mitochondrial chaperones and oxidative regulators, including Hspa1a/b, Hsp90aa1, and Gpx1, were consistently suppressed. Complementary volcano plots further highlighted the strong upregulation of lipid pathway drivers (Apoa1, Elovl4) and downregulation of enzymes linked to redox balance and mitochondrial integrity (Aifm3, Tspo2, Acoxl, Alox15, Mgst2). These combined analyses point to a transcriptional program characterized by enhanced lipid remodeling coupled with compromised mitochondrial function, reinforcing the pathway-level disruptions underlying OA-induced cardiac pathology (Supplementary Fig. 1)

These findings suggest that cytoskeletal remodeling is a shared hallmark of OA-induced heart failure, but the underlying mechanisms differ by sex. Females favor sarcomeric stiffening and pro-inflammatory activation, aligning with HFpEF-like progression, whereas males exhibit HFmrEF-like disease progression.

Transcriptomic profiling of left ventricular tissue revealed distinct and overlapping gene expression patterns in HFpEF and HFmrEF. Although most differentially expressed genes (DEGs) were unique to each condition, we identified 19 DEGs that were consistently altered across both phenotypes **(Figure 5).** These include genes involved in mitochondrial metabolism (Alas2), cell–cell adhesion and cardiac structural remodeling (Cdh2), transcriptional regulation (E2f23), RNA stability and translational control (Fam46c), and noncoding RNA processing (Snord13). The convergence of these genes suggests a shared pathogenic signature, despite the clinical and molecular divergence between HFpEF and HFmrEF. This shared core may reflect the fundamental pathways driving adverse ventricular remodeling and may represent promising targets for therapeutic strategies that could extend across the heart failure spectrum.

**Figure 5.**
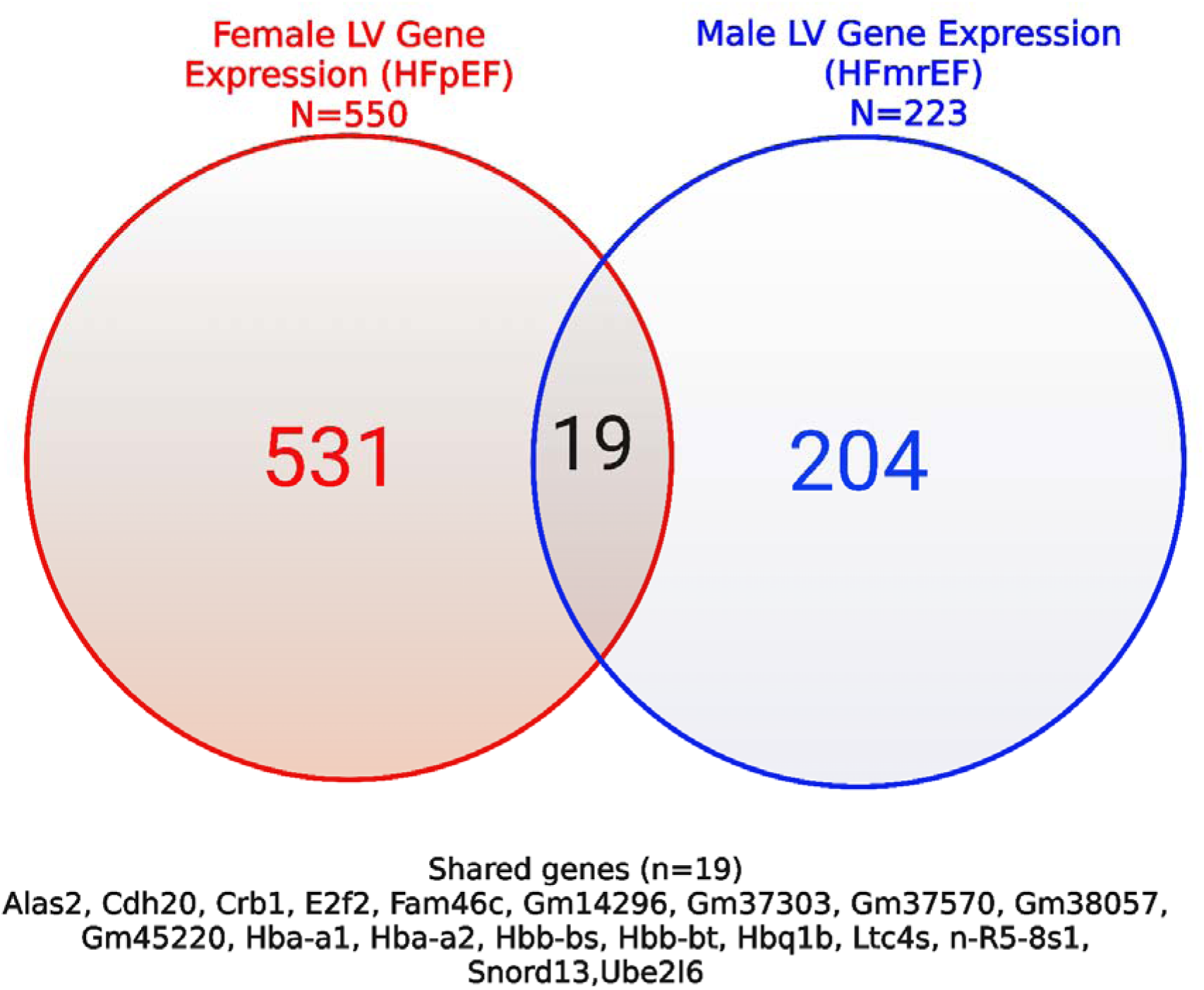
Overlap of differentially expressed genes between HFpEF and HFmrEF. The Venn diagram shows the distribution of differentially expressed genes (DEGs) in left ventricular tissue from HFpEF and HFmrEF groups. The HFpEF circle represents genes unique to heart failure with preserved ejection fraction, while the HFmrEF circle shows genes unique to heart failure with mid-range ejection fraction. The overlapping region, containing 19 genes, indicates a shared molecular signature suggesting convergent pathological mechanisms. The numbers within each section show the count of unique or shared DEGs, with the total gene count above the diagram.

### Gene Ontology (GO) enrichment analysis revealed distinct sex-specific transcriptional signatures following OA-induced heart failure

To further contextualize these transcriptional differences, we performed Gene Ontology enrichment analysis of the differentially expressed genes identified in **Figure 6**. This analysis enabled us to move beyond individual gene-level changes and evaluate the broader biological processes, molecular functions, and cellular components driving sex-specific heart failure. **In females (6A-C), Biological Processes (BP)** were enriched for *lymphocyte-mediated immunity*, *adaptive immune response*, *immunoglobulin-mediated immune response*, and *B cell-mediated immunity*. **Molecular Functions (MF)** were dominated by *cytokine activity*, *calcium channel activity*, and *oxidoreductase activity acting on peroxidases*. **Cellular Components (CC)** showed enrichment for the *collagen-containing extracellular matrix*, *cytoplasmic region*, and *plasma membrane-bound cell projections***. In males (6E-F),** Biological Processes (BP) were enriched for myeloid cell homeostasis, erythrocyte differentiation, erythrocyte homeostasis, and the hydrogen peroxide metabolic process. Molecular Functions (MF) included antioxidant activity, peroxidase activity, hemoglobin binding, and oxygen carrier activity. Cellular Components (CC) were enriched for the basolateral plasma membrane, external side of the plasma membrane, and other immune-related genes.

**Figure 6.**
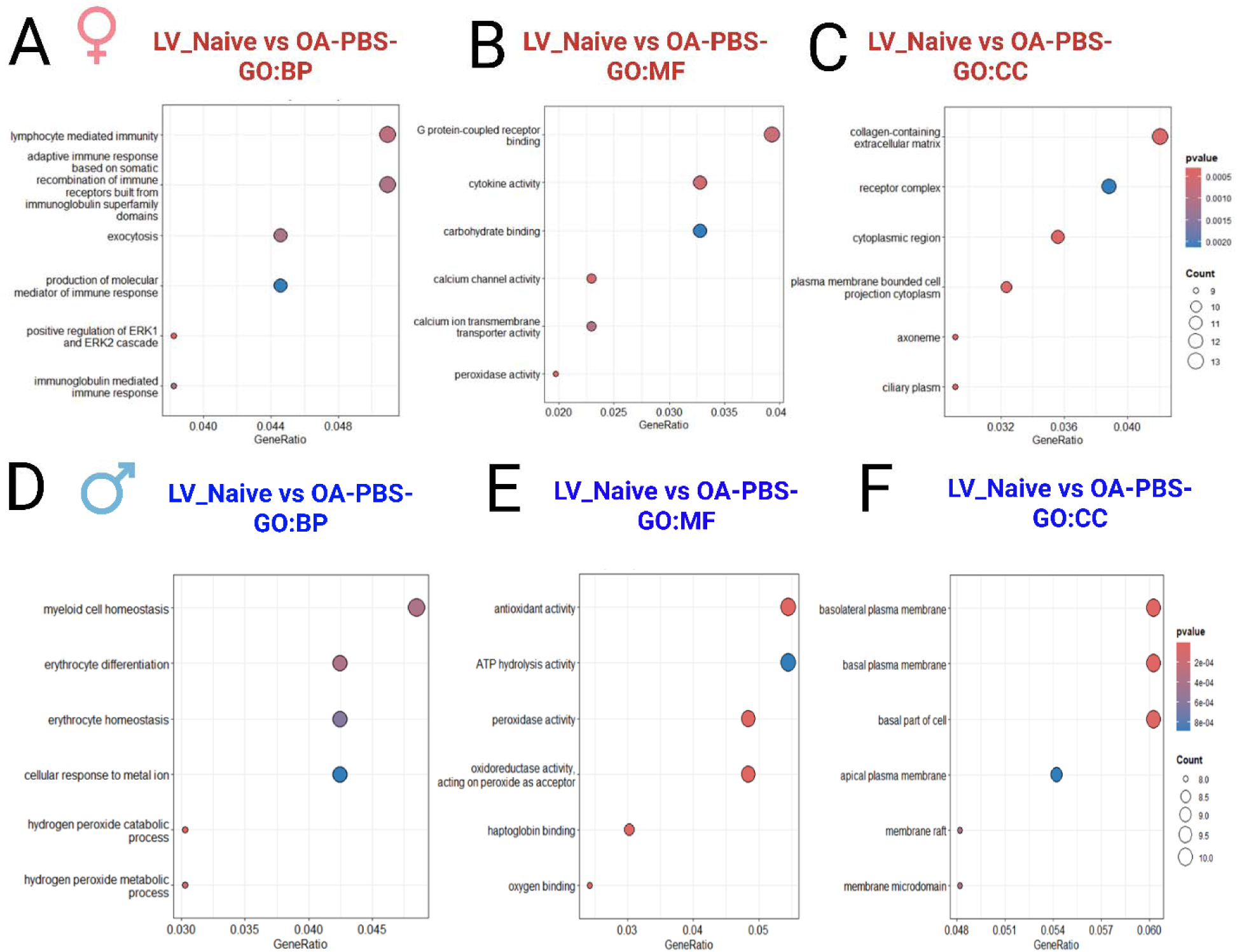
Gene Ontology enrichment analysis of differentially expressed genes in male and female following OA-induced heart failure. (A–C) In female left ventricular transcriptomes, there is an enrichment of pathways related to Biological Process (BP), Molecular Function (MF), and Cellular Component (CC), primarily involving extracellular matrix remodeling, collagen organization, and cytoskeletal regulation. (D–F) In contrast, male left ventricular transcriptomes show a predominance of immune and inflammatory pathways. The size of the circles reflects the number of genes in each category, while the color intensity indicates the –log10(p-value).

Similarly, GO-based cnet analysis (Supplementary Fig. 2) revealed sex-dependent remodeling signatures in the left ventricle following OA-induced heart failure. Male networks centered on Ttn, Actn2, and Ppargc1a reflected cytoskeletal and mitochondrial dysfunction, whereas female networks enriched for Col1a1, Lox, Calm1, and Il6 highlighted extracellular matrix and immune–calcium signaling pathways. These distinct transcriptional programs underscore the divergent molecular responses that may drive male and female heart failure phenotypes.

### Sex-Specific Alterations in Inflammatory, Oxidative Stress, and Structural Protein Networks in Osteoarthritis-Induced Heart Failure

Figure 7 illustrates the immunoblot analysis of left ventricular tissue, highlighting the distinct molecular changes associated with OA-induced cardiac dysfunction that varied by sex. In female subjects, OA prompted a notable increase in proteins linked to hypertrophy and fibrosis, such as ANP, BNP, and the Col1/Col3 ratio, suggesting structural cardiac changes alongside elevated expression of inflammatory pathway genes, including p38, p65, and Gal4. Furthermore, female mice showed heightened expression of genes related to hypoxic conditions and oxidative stress, namely Hif1a, 4HNE, Ero1, Sirt1, and AMPK, without any alterations in the expression of genes associated with the unfolded protein response or autophagy. In contrast, male participants with OA exhibited similar elevations in structural and inflammatory pathways (ANP, Col1/Col3, p38, and p65), coupled with reductions in proteins related to the unfolded protein response (Ero1, Prdx4, and PDI) and an increase in autophagic proteins (SQSTM1, LC3B, M6PR, Ubiquitin). Notably, female hearts demonstrated a significant reduction in cleaved caspase 3, whereas male hearts showed a significant increase, highlighting a crucial sex-specific difference in apoptotic activity.

**Figure 7.**
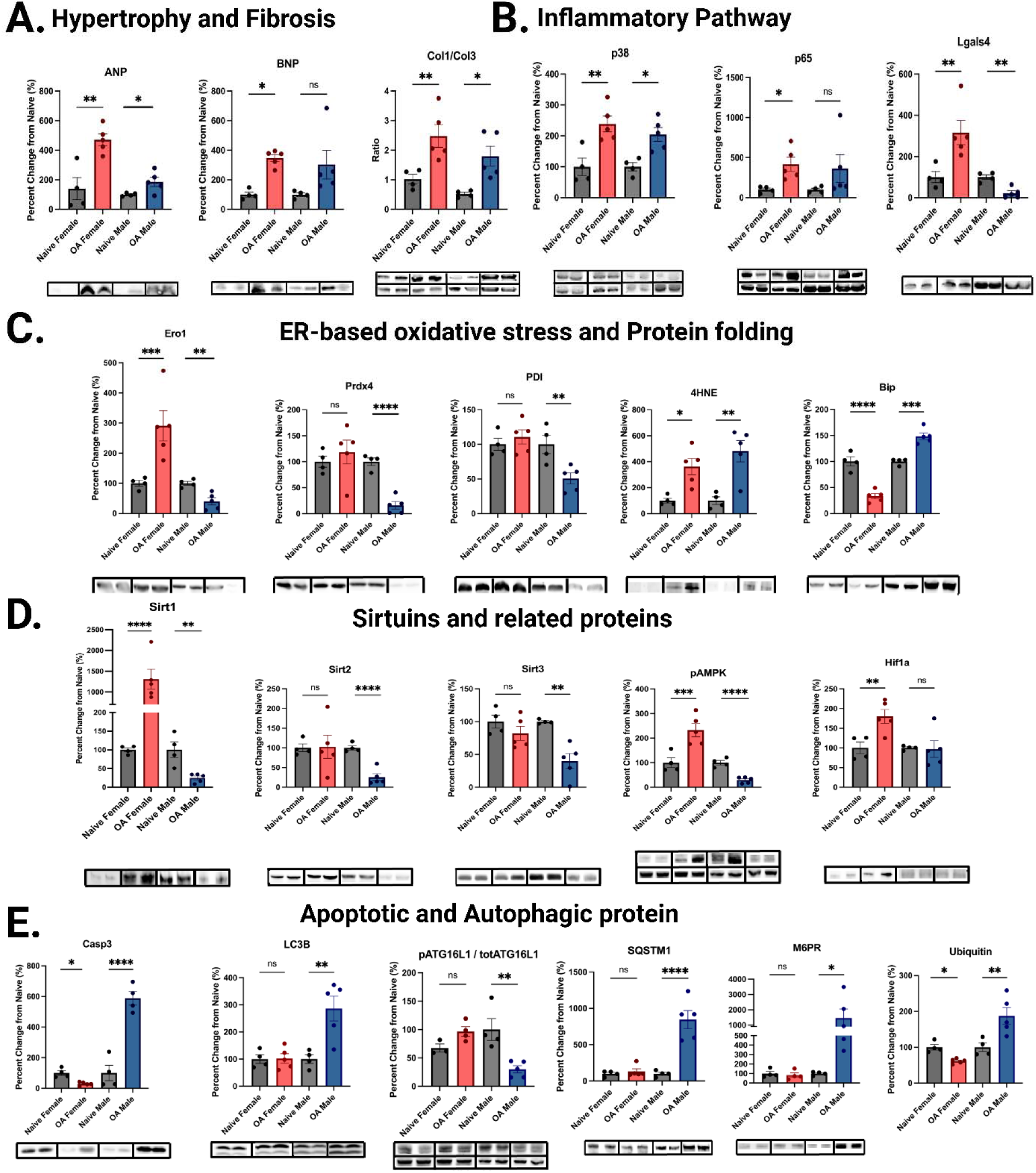
Representative immunoblot analyses of left ventricular tissue from naïve females, OA females, naïve males, and OA males. Protein abundance was quantified relative to total protein loading, with phosphorylated forms presented above corresponding to total protein. (A) Cardiac structural and functional biomarkers. (B) Inflammation-associated proteins. (C) ER-based oxidative stress and protein folding-related proteins. (D) Sirtuins and associated regulatory proteins. (E) Apoptotic and autophagic signaling proteins. Representative blots for each group are displayed below the corresponding quantification. One-way ANOVA determined statistical significance with post hoc testing; *P < 0.05, **P < 0.01, ***P < 0.001, ****P < 0.0001.

Quantitative analyses demonstrated elevated phosphorylation of signaling proteins in OA males, consistent with the activation of inflammatory and cell death pathways. In women with OA, increased expression of sirtuins and other regulatory proteins associated with metabolic and stress resilience has been observed. Although these data were generated from separate analyses, the findings suggest sex-dependent molecular responses to OA that may contribute to the divergent pathophysiological outcomes.

## Discussion

Heart failure (HF) has transitioned from being viewed as a disease of isolated myocardial injury to a systemic syndrome driven by persistent inflammation and metabolic stress. Among the less traditional contributors, osteoarthritis (OA), the most prevalent degenerative joint disease, has emerged as a significant yet under-recognized risk factor for cardiovascular pathology. Large-scale clinical studies have consistently shown that patients with OA face up to a three-fold higher risk of developing HF, even after accounting for age, obesity, and reduced mobility (13–15). This elevated risk reflects a shared pathophysiological axis linking low-grade chronic inflammation, endothelial dysfunction, and metabolic imbalance, which accelerates cardiac remodeling (7, 8). Mounting evidence indicates that inflammation plays a central role in the initiation and progression of HF with preserved ejection fraction (HFpEF), a condition that predominantly affects postmenopausal women(10, 23–25). Chronic inflammatory disorders, such as OA, autoimmune diseases, metabolic syndrome, and aging-associated inflammaging, have all been identified as key risk factors for HFpEF(7, 8). However, clinical trials targeting single inflammatory pathways, such as IL-1β inhibition, have largely failed to improve HFpEF outcomes (26), underscoring the heterogeneous and sex-dependent nature of the inflammation underlying this syndrome. Inflammation-driven cardiac dysfunction is multifactorial and context-dependent, with distinct immune, metabolic, and vascular components contributing to divergent heart failure trajectories in men and women.

In this study, we developed the first preclinical model of osteoarthritis-induced heart failure (OA-HF) to establish OA as a direct causal driver of cardiac dysfunction. Surgical destabilization of the medial meniscus triggers chronic systemic inflammation and produces distinct sex-specific cardiac phenotypes: females exhibit HFpEF-like diastolic dysfunction, whereas males develop HFmrEF-like systolic impairment. Notably, HFmrEF represents a dynamic and transitional stage that can evolve toward heart failure with reduced ejection fraction (HFrEF), capturing a continuum of inflammatory and structural decline(1, 11). This gender-specific pattern reflects clinical observations, where women are more likely to develop HFpEF and men are at a higher risk of progressive systolic failure (12).

### Sex-Specific Echocardiographic and Functional Characterization of OA-Induced Heart Failure

Longitudinal echocardiography has revealed distinct sex-dependent trajectories of cardiac remodeling that parallel the clinical continuum of human heart failure. Representative M-mode, pulse-wave Doppler (E/A ratio), and tissue Doppler (E/e′) recordings demonstrated preserved systolic function with impaired diastolic relaxation in females—consistent with heart failure with preserved ejection fraction (HFpEF)— and progressive systolic decline in males, characteristic of heart failure with mid-range ejection fraction (HFmrEF), a dynamic transitional phase toward heart failure with reduced ejection fraction (HFrEF) (Fig. 2A–B). In females, the ejection fraction remained preserved through 16 weeks post-OA, while diastolic indices exhibited a temporal progression marked by early E/A reduction (8–10 weeks), mid-phase pseudo-normalization (10–12 weeks), and late-stage elevation (13–16 weeks), accompanied by increased E/e′ and prolonged isovolumetric relaxation time (IVRT). These temporal dynamics recapitulate the established echocardiographic evolution of human HFpEF, in which E/A reversal, elevated E/e′, and extended IVRT signal increase ventricular stiffness and impair relaxation (27, 28). In contrast, males displayed an early decline in systolic performance beginning at week 10, with concurrent increases in E/e′ and prolonged isovolumetric contraction time (IVCT), indicating delayed contractility and elevated filling pressures, which are hallmarks of the HFmrEF/HFrEF spectrum (2, 15).

Notably, our data revealed that male OA mice remained within the HFmrEF window rather than progressing to full HFrEF by 16 weeks, likely reflecting the role of systemic inflammation as a major contributor to OA-induced cardiac dysfunction. OA-driven inflammatory signaling—marked by endothelial stress, immune activation, and metabolic dysregulation—appears to initiate diastolic dysfunction in females by week 8 and systolic decline in males by week 10, defining sex-specific windows of vulnerability (7, 8, 13, 16). Given that our longitudinal observations concluded at week 16, this HFmrEF phenotype likely represents an evolving transitional stage that, if extended to 20 weeks or beyond, would be expected to advance to overt HFrEF. Together, these results establish OA-induced systemic inflammation as a sufficient and sex-differentiated driver of heart failure progression.

Morphometric analyses further underscored these divergent remodeling patterns: females exhibited concentric hypertrophy, as evidenced by an increased heart weight-to-tibia length ratio (HW/TL³), whereas males displayed eccentric remodeling with chamber dilation and wall thinning characteristic of systolic failure. These findings are consistent with the established literature demonstrating that female hearts typically mount a concentric hypertrophic response and preserve systolic function, whereas male hearts more readily develop eccentric remodeling and systolic decline (29, 30).Mechanistically, these sex-specific adaptations are thought to arise from the differential activation of signaling pathways; females preferentially engage ERβ–Akt–MAPK signaling, which promotes adaptive growth and mitochondrial homeostasis, whereas males show heightened androgen- and CaMKII-driven responses that exacerbate fibrosis, apoptosis, and contractile dysfunction (29, 30).

### Transcriptomic Signatures Underlying Sex-Specific Remodeling in Osteoarthritis-Induced Heart Failure

Bulk RNA sequencing of left ventricular (LV) tissue revealed a clear sexual dimorphism in transcriptional remodeling following osteoarthritis (OA)-induced systemic inflammation, consistent with prior clinical and experimental studies (31–33). The clustering of distinctly separated female and male transcriptomes effectively recapitulated the divergent phenotypes of heart failure, specifically diastolic-preserved HFpEF in females and systolic-declining HfmrEF/HfrEF in males.

In the left ventricle of osteoarthritis (OA)-induced mice, transcriptomic profiling revealed distinct sex-dependent molecular remodeling programs (Fig. 4–6, Table 1). In female hearts, genes associated with extracellular matrix (ECM) organization, cytoskeletal remodeling, and calcium handling, including Col1a1, Col3a1, Ttn, Neb, and Kcnab2, were significantly upregulated, with enrichment in focal adhesion, collagen fibril assembly, and calcium ion–binding pathways. These transcriptional signatures correspond to concentric hypertrophy and diastolic stiffness, hallmarks of heart failure with preserved ejection fraction (HFpEF) (Fig. 4). This ECM-dominant profile aligns closely with human LV transcriptomes from HFpEF biopsies, which demonstrate robust collagen cross-linking, titin hypophosphorylation, and integrin-mediated adhesion remodeling in the absence of classical inflammatory activation (31, 34). In contrast, male LV transcriptomes were enriched for inflammatory and immune activation networks, characterized by the upregulation of Tnf, Ccl2, Cxcl10, Nfkb1, and Stat3, and Gene Ontology enrichment for cytokine-mediated signaling, leukocyte chemotaxis, and interferon-γ responses (Fig. 5A–C). This immune-driven remodeling mirrors the molecular landscape of the human HFrEF myocardium, in which macrophage recruitment, STAT4 activation, and NFκB-driven cytokine cascades precipitate systolic dysfunction and eccentric remodeling (32, 33, 35).

These transcriptomic signatures fit within an emerging cardio-immunologic framework distinguishing “outside-in” systemic inflammation in HFpEF from “inside-out” myocardial inflammation in HFrEF (Supplementary Fig. 1 and 2). In HFpEF, systemic low-grade inflammationarising from comorbidities such as obesity, diabetes, or OAactivates circulating immune cells and the vascular endothelium, leading to paracrine fibroblast stimulation, ECM deposition, and progressive diastolic dysfunction (33). Consistent with this, transcriptomic analyses of peripheral blood mononuclear cells (PBMCs) from patients with HFpEF reveal B-cell enrichment, cytokine receptor signaling, and metabolic reprogramming, whereas LV biopsies show minimal induction of canonical inflammatory transcripts. In contrast, HFrEF follows an “inside-out” paradigm, initiated by primary myocardial injury (e.g., ischemia, pressure overload) that releases damage-associated molecular patterns (DAMPs), recruits CCR2⁺ monocytes/macrophages, and activates Th1-polarized T cells within the myocardium (36).

Across both sexes, the coordinated downregulation of mitochondrial and lipid metabolic genes (Cpt1b, Ndufs2, Ppargc1a) indicated impaired oxidative metabolism and bioenergetic inefficiency, which are shared hallmarks of chronic inflammation-induced cardiac dysfunction (Fig. 6). This overlap in TNF–NFκB signaling, immune-metabolic stress, and mitochondrial dysfunction underscores a unifying pathogenic axis that bridges the female ECM-fibrotic (HFpEF-like) and male cytokine-driven (HFrEF-like) remodeling trajectories. Collectively, these findings establish the OA-induced model as a translationally relevant platform for dissecting sex-specific inflammatory remodeling in heart failure and highlight convergent therapeutic targets within the mitochondrial and TNF–NFκB signaling pathways.

### Sex-Dependent Molecular Remodeling in the Left Ventricle Revealed by Immunoblot Analysis Following Osteoarthritis-Induced Cardiac Dysfunction

Immunoblot analyses revealed distinct sex-dependent molecular remodeling in the left ventricle following osteoarthritis (OA)-induced cardiac dysfunction. OA triggered coordinated changes in proteins regulating hypertrophy, fibrosis, inflammation, oxidative stress, and apoptosis, underscoring the divergent mechanisms of cardiac injury between the sexes (Fig. 7). Consistent with prior studies, female hearts exhibited a fibrotic and hypertrophic profile characteristic of HFpEF, whereas male hearts displayed greater inflammatory and apoptotic activation typical of HFrEF(12, 29). These findings highlight biological sex as a key determinant of molecular adaptation and disease trajectory in inflammation-driven cardiac remodeling.

### Hypertrophy and Fibrosis-Related Proteins

Both sexes showed increased atrial natriuretic peptide (ANP) and brain natriuretic peptide (BNP) levels and elevated collagen I/III ratios, which are hallmarks of pressure overload and myocardial fibrosis (Ye et al., 2023). These changes reflect maladaptive remodeling with extracellular matrix expansion and diastolic stiffening, particularly in females, consistent with HFpEF-associated fibrotic remodeling(12). Transcriptomic analyses of the human HFpEF myocardium similarly demonstrate the upregulation of collagen synthesis and matrix-regulatory genes without overt inflammatory activation(31). Collectively, chronic mechanical and cytokine stress appears to drive a self-sustaining fibrotic response through TGF-β, CTGF, and collagen signaling, which is most evident in estrogen-rich female hearts (36).

### Inflammatory Signaling Pathways

OA induced robust activation of inflammatory signaling, as evidenced by the increased phosphorylation of p38 MAPK, NF-κB p65, and Gal4, which are key mediators of cytokine-driven remodeling (36). This inflammatory surge promotes maladaptive hypertrophy and contractile impairment, aligning with the known TNF-α– and IL-1β–dependent pathways in heart failure (36). Male hearts exhibited stronger kinase phosphorylation, consistent with heightened immune cell infiltration and greater proinflammatory remodeling, whereas female hearts displayed concurrent upregulation of HIF-1α and 4-HNE, indicating an adaptive redox response to metabolic stress rather than overt inflammation. These findings parallel those of Alcaide et al. who observed that HFpEF involves low-grade endothelial and fibroblast activation, in contrast to the more aggressive inflammatory milieu observed in HFrEF (36).

### Endoplasmic Reticulum Stress and Oxidative Homeostasis

Distinct sex-dependent differences were observed in oxidative and endoplasmic reticulum (ER) stress responses. Female hearts exhibited higher levels of Ero1, Sirt1, and AMPK, suggesting enhanced redox regulation and metabolic adaptability, whereas male hearts showed reduced expression of ER chaperones, such as Ero1, Prdx4, and PDI, indicative of impaired protein folding and heightened oxidative burden. These disparities mirror prior evidence that the myocardium of males displays greater oxidative susceptibility due to lower antioxidant enzyme activity and mitochondrial resilience (37). Activation of the Sirt1–AMPK axis in females likely preserves mitochondrial function and energy efficiency by maintaining NAD⁺ redox balance and supporting adaptive biogenesis (38). Together, these mechanisms underscore a sexually dimorphic redox landscape in which females exhibit stress tolerance through metabolic flexibility, whereas males are predisposed to proteostatic collapse and oxidative injury.

### Sirtuin- and Metabolism-Linked Pathways

Enhanced activation of Sirt1 and AMPK in females with OA reflects a coordinated shift toward metabolic resilience and efficient energy utilization. Sirtuins regulate mitochondrial function and longevity through PGC-1α–dependent deacetylation, promoting oxidative phosphorylation and limiting reactive oxygen species generation (39). In contrast, male hearts displayed attenuated Sirt1 signaling and greater kinase phosphorylation, indicative of metabolic inflexibility and energy depletion. Such differences align with reports that female myocardium maintains superior mitochondrial efficiency and substrate flexibility, favoring fatty acid oxidation under stress, whereas males rely more heavily on glycolytic flux, predisposing them to energetic failure (40). These data suggest that the preservation of the Sirt1–AMPK axis in females confers metabolic adaptability, whereas its suppression in males contributes to the transition from compensated remodeling to contractile dysfunction.

### Apoptosis and Autophagy

Sex-specific differences were also evident in the pathways governing apoptosis and autophagy. Female hearts exhibit reduced levels of cleaved caspase-3, reflecting estrogen-mediated cytoprotection and attenuation of apoptotic signaling (41). In contrast, male hearts displayed marked upregulation of caspase-3, LC3B, and SQSTM1, indicating the concurrent activation of apoptotic and autophagic mechanisms. While autophagy initially supports cellular homeostasis by clearing damaged proteins, chronic activation under oxidative or ER stress can exacerbate cell death and ventricular dysfunction (42). This dual pattern suggests that females rely on adaptive autophagy and survival signaling to preserve cardiomyocyte integrity, whereas males experience maladaptive proteostatic stress that accelerates the structural deterioration.

Together, these findings delineate a unified model in which osteoarthritis (OA) induces systemic inflammation and metabolic stress, driving sex-dependent cardiac remodeling. Female hearts engage adaptive signaling networks that promote redox balance, metabolic flexibility, and extracellular matrix stabilization, yielding a fibrotic and HFpEF-like phenotype. In contrast, male hearts exhibit energetic collapse, ER dysfunction, and apoptotic remodeling, which are characteristic of HfrEF(12, 36). These dimorphic responses mirror hormonal influences, with estrogen enhancing mitochondrial function and SIRT1–AMPK signaling, while testosterone amplifies inflammatory kinase activation and cell death cascades (38, 40). Collectively, these data position OA as a systemic inflammatory disorder capable of precipitating divergent molecular trajectories of heart failure. Understanding these sex-specific proteomic and metabolic adaptations provides a mechanistic foundation for developing precision therapies that target oxidative stress, AP, and inflammation in a sex-specific manner.

## Conclusion

This study established osteoarthritis (OA) as a systemic inflammatory driver of sex-specific heart failure (HF) using a novel murine model. Chronic inflammation induced by destabilization of the medial meniscus (DMM) produces distinct cardiac phenotypes: females develop diastolic dysfunction with preserved ejection fraction (HFpEF), whereas males exhibit progressive systolic impairment consistent with HFmrEF/HFrEF. These trajectories mirror the clinical patterns of HF and underscore biological sex as a key determinant of inflammatory cardiac remodeling. Transcriptomic and proteomic analyses revealed shared mitochondrial and metabolic dysfunction but divergent molecular adaptations between the sexes. Female hearts demonstrated extracellular matrix and Sirt1–AMPK activation consistent with fibrotic remodeling and metabolic resilience, whereas male hearts displayed heightened p38 MAPK and NF-κB signaling, coupled with increased autophagy and apoptosis. Collectively, these data define inflammation as a unifying mechanism linking musculoskeletal degeneration and cardiac dysfunction. By integrating joint degeneration and cardiac remodeling within a single experimental framework, the OA-induced HF model provides a physiologically relevant and translational platform for dissecting immune–metabolic interactions and identifying sex-specific therapeutic targets for inflammation-driven heart failure.

## Summary

**Figure 8.**
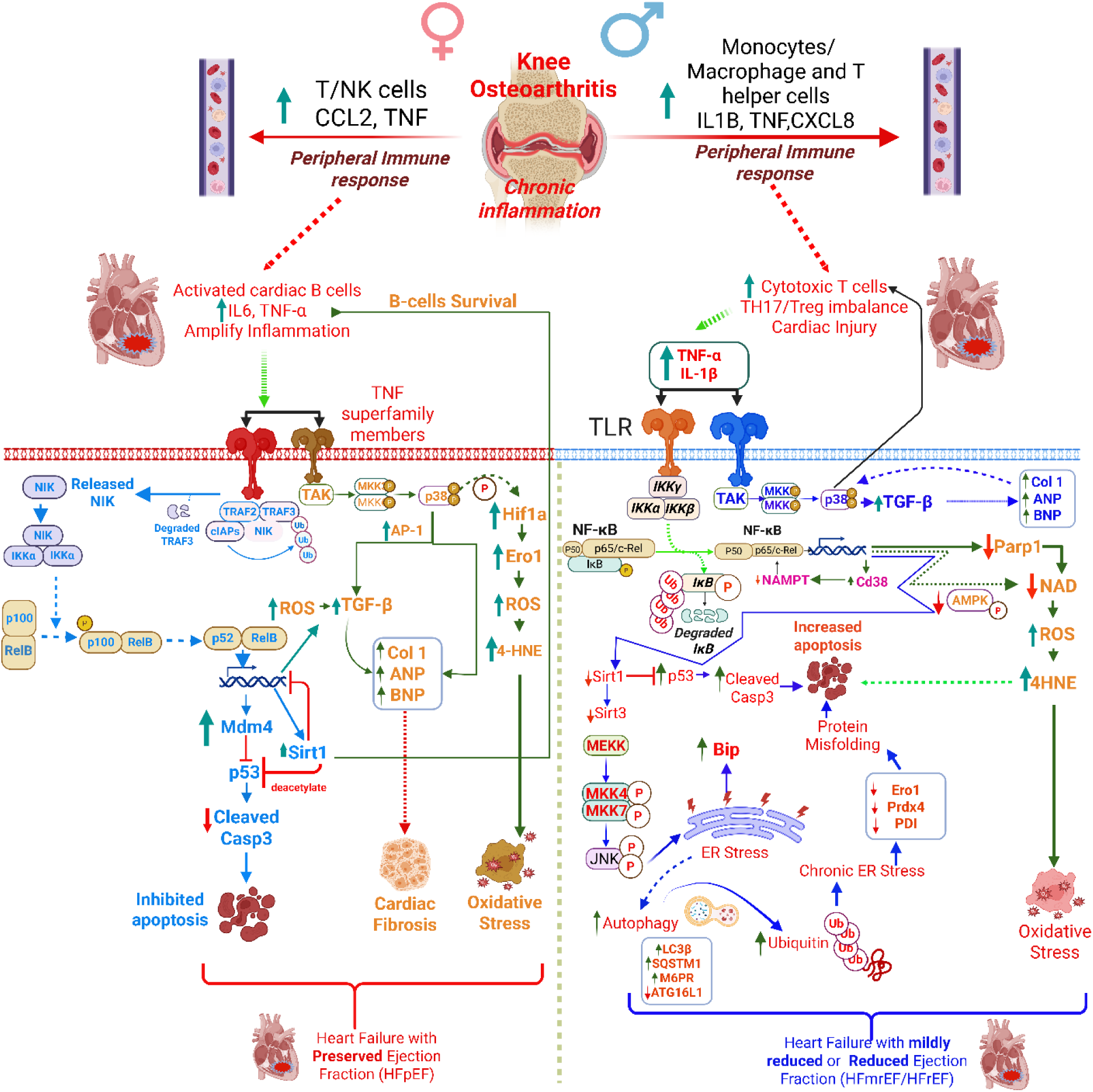
Proposed mechanism of osteoarthritis (OA)-induced, sex-specific heart failure phenotypes. Knee osteoarthritis (OA) elicits chronic joint inflammation, which triggers systemic immune activation and sustained release of proinflammatory mediators, such as TNF and CCL2. These circulating cytokines and activated immune cells impair vascular homeostasis, induce endothelial dysfunction, and promote metabolic stress in peripheral organs, including the heart. In females (left, blue), this represents an “outside-in” inflammatory mechanism, in which systemic immunometabolic activation precedes cardiac dysfunction. Circulating T and NK cells display enhanced cytokine signaling and oxidative metabolism, upregulating *Sirt1*, *Prkaa1 (AMPK)*, *Nppa*, and *Nppb* to support metabolic resilience and diastolic compliance in HFpEF. Despite minimal local inflammation, persistent low-grade systemic activation leads to endothelial impairment, fibrosis, and concentric hypertrophy, which are hallmarks of heart failure with preserved ejection fraction (HFpEF). In males (right, red), OA promotes an “inside-out” mechanism initiated by cardiomyocyte stress and necrosis, resulting in robust NF-κB and p38 MAPK activation. This triggers oxidative stress, autophagy (*Lc3b*), and apoptosis (*Casp3*), accompanied by increased *Tnf*, *Il6*, and *Il1b* expression, which together drive monocyte/macrophage recruitment, B cell activation, and Th17/Treg imbalance. The ensuing inflammation amplifies tissue remodeling, fibrosis, and eccentric hypertrophy, which are characteristic of heart failure with reduced ejection fraction (HFrEF) and its transitional form (HFmrEF). Collectively, this model illustrates how localized OA inflammation propagates systemic immune dysregulation that manifests as distinct cardiac phenotypes—immunometabolic, fibrotic HFpEF in females versus injury-driven, proinflammatory HFrEF in males—underscoring the need for sex- and phenotype-specific therapeutic strategies for HF. ***Selected gene abbreviations: CCL2 (C-C Motif Chemokine Ligand 2), TNF (Tumor Necrosis Factor), IL1B (Interleukin 1 Beta), CXCL8 (C-X-C Motif Chemokine Ligand 8 / IL-8), Nfkb1 (Nuclear Factor Kappa B Subunit 1), Mapk14 (p38 MAPK), Sirt1 (Sirtuin 1), Prkaa1 (AMP-Activated Protein Kinase α1), Lc3b (Microtubule-Associated Protein 1 Light Chain 3β), Casp3 (Caspase-3), Rock1 (Rho-Associated Coiled-Coil Kinase 1), Gabra2 (Gamma-Aminobutyric Acid A Receptor Subunit Alpha 2)*.**

## Abbreviations

OA: Osteoarthritis
DMM: Destabilization of the Medial Meniscus
HF: Heart Failure
HFpEF: Heart Failure with Preserved Ejection Fraction
HFmrEF: Heart Failure with Mildly Reduced Ejection Fraction
HFrEF: Heart Failure with Reduced Ejection Fraction
ECM: Extracellular Matrix
LV: Left Ventricle
EF: Ejection Fraction
IVRT: Isovolumetric Relaxation Time
IVCT: Isovolumetric Contraction Time
RNA-seq: RNA Sequencing

## Data availability statements

Data available upon request; contact John R. Bethea and Pranav Prasoon (email ID)

## Author Contributions

**Conceptualization:** Pranav Prasoon and John R. Bethea, **DMM Surgery and Von Frey testing:** Pranav Prasoon, **WB and Echo:** Kelly Tammen, **RNAseq sample preparation and analysis:** Aravind Meyyappan, Kelly Tammen and Pranav Prasoon; **Histology:** Manushri Dalvi. **Original draft preparation: Pranav** Prasoon; **Schematic diagrams:** Kelly Tammen and Pranav Prasoon; **tables created:** Pranav Prasoon and Kelly Tammen; **Writing, review, and editing:** all authors.

