## Supplementary Figure 1-3 and Table 1-2 for "Sex-Specific Characterization of a Novel Osteoarthritis-Induced Heart Failure Model in Mice": Supplematary Fig 2.docx

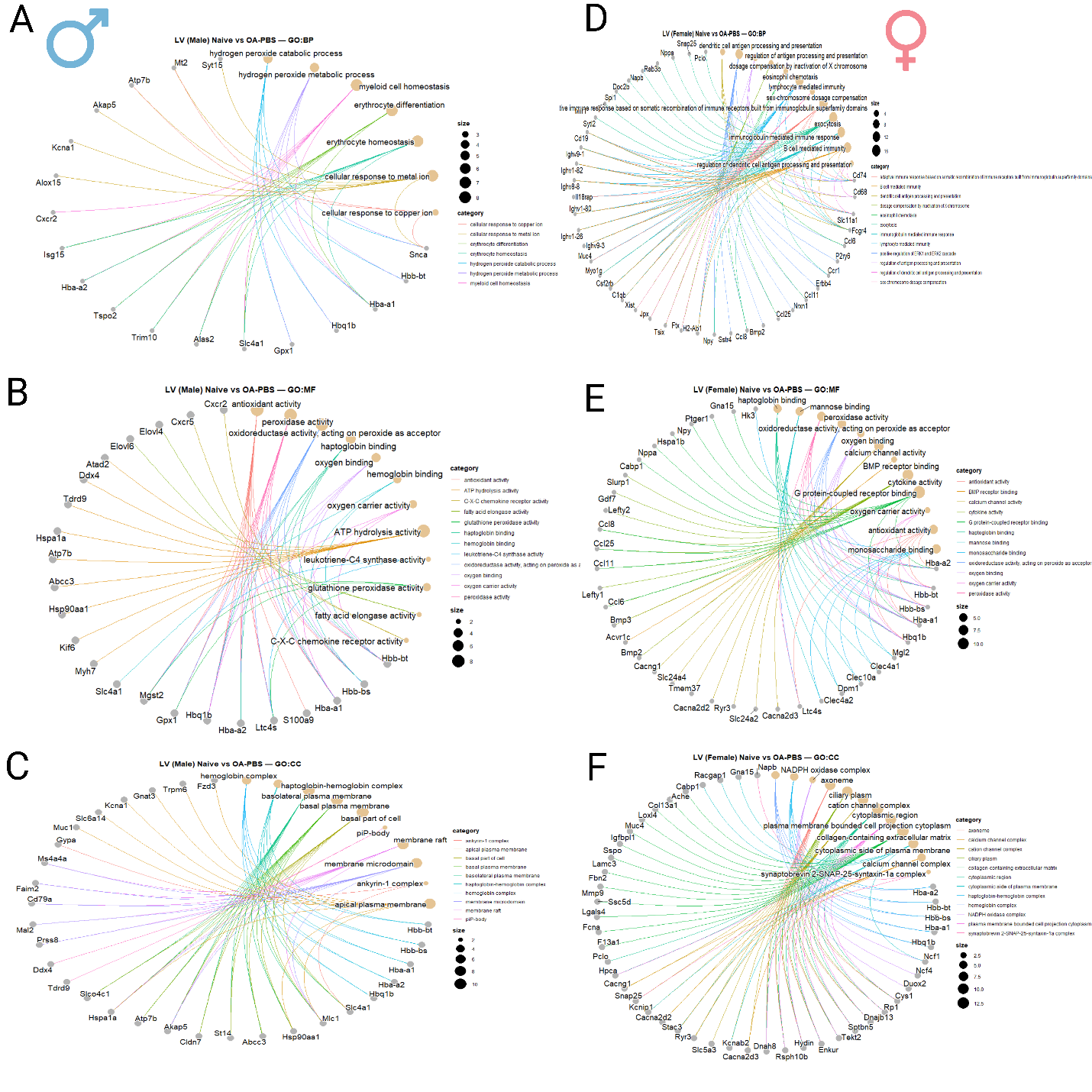

Supplementary Figure 2 | **Cnet plots of Gene Ontology enrichment in the left ventricle of male and female mice following osteoarthritis-induced heart failure.**

(A–C) Gene–concept network (cnet) plots of enriched biological processes (BP), molecular functions (MF), and cellular components (CC) in male left ventricle tissue following osteoarthritis (OA)–induced heart failure.

(D–F) Corresponding GO cnet plots for female left ventricle tissue.

Nodes represent individual genes (circles) or GO terms (diamonds), with connecting edges denoting gene-to-term associations. Node size reflects the number of associated genes, and color intensity corresponds to enrichment significance (–log₁₀ p-value). Males exhibited enrichment of pathways related to cytoskeletal organization, mitochondrial metabolism, and lipid catabolism, whereas females displayed activation of extracellular matrix organization, calcium signaling, and immune regulatory pathways. These sex-specific GO networks reveal divergent molecular programs underlying cardiac remodeling after OA-induced injury.
