## Supplementary Figure 1-3 and Table 1-2 for "Sex-Specific Characterization of a Novel Osteoarthritis-Induced Heart Failure Model in Mice": Supplementary Figure 1.docx

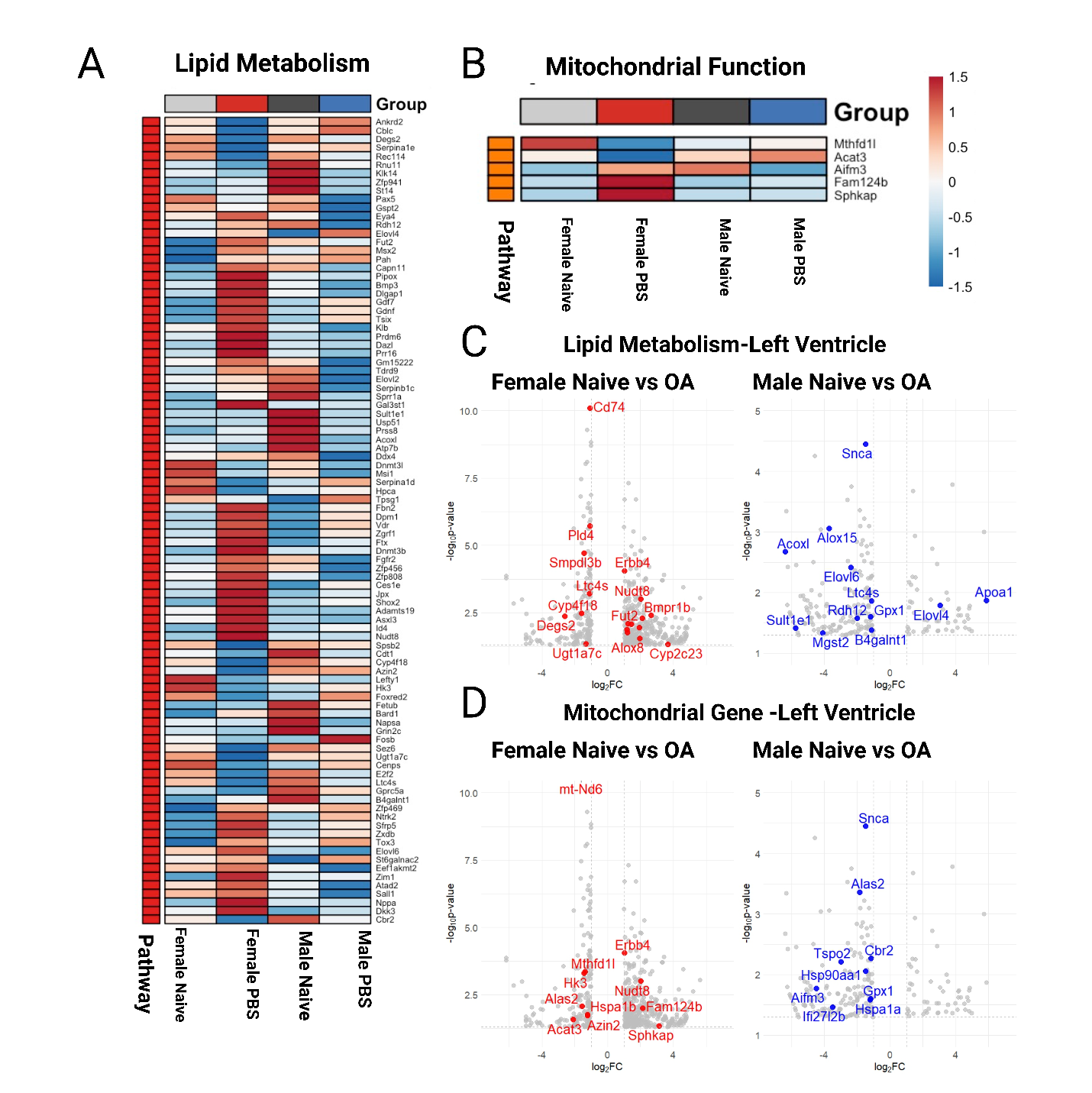

**Supplementary Figure 1. Transcriptomic analysis of left ventricular tissue reveals pathway-level dysregulation in lipid metabolism and mitochondrial function following OA-induced heart failure.**
(A–B) Heatmaps illustrate differential expression of all significant genes annotated to **lipid metabolism** (A) and **mitochondrial function** (B), clustered by normalized expression values from bulk RNA sequencing. Patterns highlight distinct molecular signatures across naïve, OA-PBS, and TNFR2 agonist–treated groups in both sexes. (C–D) Volcano plots display significantly upregulated and downregulated genes within the **lipid metabolism** (C) and **mitochondrial** (D) gene sets, emphasizing pathways linked to altered bioenergetics, metabolic remodeling, and mitochondrial health.
