## Supplementary Figure 1-3 and Table 1-2 for "Sex-Specific Characterization of a Novel Osteoarthritis-Induced Heart Failure Model in Mice": Supplementary Figure 3.docx

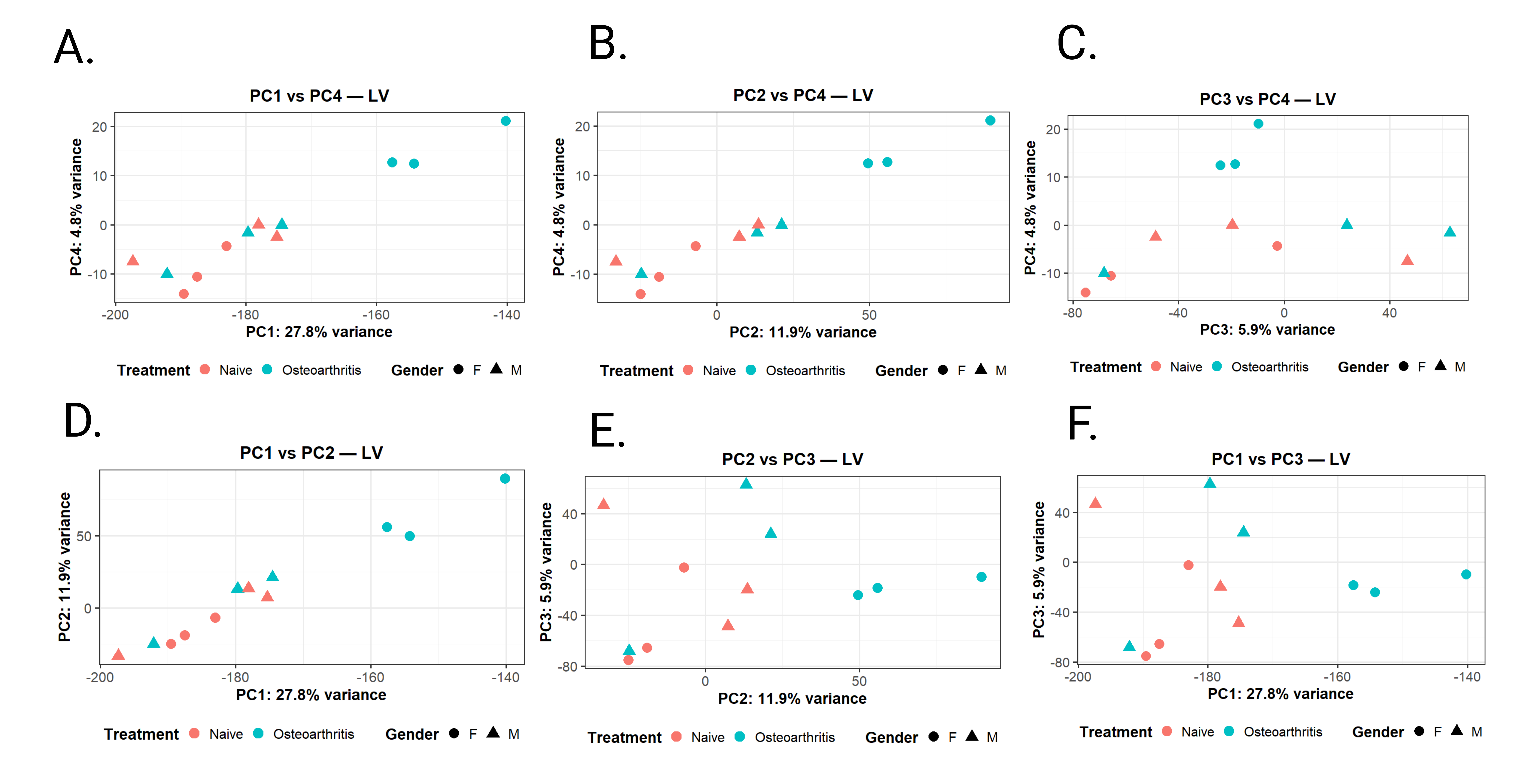

Supplementary Figure 3: **Distinct transcriptional clustering of naïve and osteoarthritic left ventricles.**

**Principal component analysis (PCA)** of normalized RNA-sequencing data was performed to assess overall transcriptional variation in left ventricular tissue from naïve and osteoarthritis (OA) mice. Each point represents an individual sample, with color indicating treatment group (red, naïve; blue, OA) and shape denoting sex (circle, female; triangle, male). Shown are pairwise comparisons of the first four principal components (PCs), which together explain the majority of total variance across samples: **A**, PC1 vs. PC4; **B**, PC2 vs. PC4; **C**, PC3 vs. PC4; **D**, PC1 vs. PC2; **E**, PC2 vs. PC3; **F**, PC1 vs. PC3. The percentage of variance explained by each component is indicated on the corresponding axes. Separation of naïve and OA samples, along with partial sex-specific clustering, highlights disease- and sex-dependent remodeling of left ventricular transcriptional profiles.
