## Supplementary Figure 1-3 and Table 1-2 for "Sex-Specific Characterization of a Novel Osteoarthritis-Induced Heart Failure Model in Mice": Table 1.docx

**Table 1: Significant Upregulated and Downregulated gene in heart left ventricle (LV) of male and female following Novel Osteoarthritis-induced Heart Failure**

| Female | | Male | |
| --- | --- | --- | --- |
| Gene Symbol | **Log2FC** | **Gene Symbol** | **Log2FC** |
| Upregulated Gene | | **Upregulated Gene** | |
| Npy | 3.724344762 | Apoa1 | 5.867932686 |
| Bmpr1b | 2.640685411 | Nkg7 | 4.17215957 |
| Nppa | 1.852125394 | Cd96 | 3.43119995 |
| Mas1 | 1.375059316 | **Downregulated Gene** | |
| Bmp2 | 1.250570795 | Gpx1 | -1.17654102 |
| Ccl25 | 1.096819148 | S100a9 | -1.34944444 |
| Npy | 3.724344762 | Snca | -1.488085739 |
| Downregulated Gene | | Fanca | -1.512822036 |
| Themis2 | -1.006134457 | Ifi206 | -1.927011795 |
| Ccl6 | -1.033292395 | Cxcr2 | -2.889608531 |
| Il10ra | -1.048298375 | Alox15 | -3.713626773 |
| Cd68 | -1.068985937 | Tnfsf18 | -4.781439027 |
| Bpgm | -1.074345404 | Rab44 | -5.214935477 |
| Slc11a1 | -1.074509333 | Elf3 | -5.635589102 |
| Pld4 | -1.077841498 |  |  |
| Pstpip1 | -1.088503174 |  |  |
| Ccl11 | -1.101724435 |  |  |
| Ccl8 | -1.182650567 |  |  |
| Ncf1 | -1.19483589 |  |  |
| Clec10a | -1.215553505 |  |  |
| Ly86 | -1.216740179 |  |  |
| Pik3ap1 | -1.218579976 |  |  |
| Ptger1 | -1.405252934 |  |  |
| Smpdl3b | -1.442841061 |  |  |
| Ccr1 | -1.493310161 |  |  |
| Fcgr4 | -1.507737631 |  |  |
| Il18rap | -1.815030253 |  |  |
| Cebpb | -1.613891546 |  |  |

**Inflammatory Genes**

**Immune Genes**

| Female | | Male | |
| --- | --- | --- | --- |
| Gene Symbol | **Log2FC** | **Gene Symbol** | **Log2FC** |
| Upregulated Gene | | **Upregulated Gene** | |
| Ighv1-82 | 4.458475099 | **Apoa1** | 5.867932686 |
| Fcamr | 4.315980858 | **Il12a** | 4.179161935 |
| Ighv1-80 | 3.940065761 | **Nkg7** | 4.17215957 |
| Cfhr2 | 3.755657177 | **Gm4841** | 3.790312294 |
| Npy | 3.724344762 | **Exo1** | 3.516626965 |
| Igkv5-43 | 3.710200285 | **Cd96** | 3.43119995 |
| Igkv15-103 | 3.582729374 | **Gm12185** | 2.564246628 |
| Ighv1-26 | 3.523936276 | **Cd3d** | 2.343627344 |
| Muc4 | 2.407977288 | **Lif** | 2.097601478 |
| Rab3b | 1.964608655 | **Downregulated Gene** | |
| Nppa | 1.852125394 | **Isg15** | -1.047543532 |
| Lgals4 | 1.273741178 | **Gpx1** | -1.17654102 |
| Ccl25 | 1.096819148 | **Cd79a** | -1.301958115 |
| Downregulated Gene | | **Igkc** | -1.317863248 |
| Cd86 | -1.00015766 | **S100a9** | -1.34944444 |
| Themis2 | -1.006134457 | **Hsp90aa1** | -1.483301564 |
| C1qb | -1.008329065 | **Snca** | -1.488085739 |
| Myo1g | -1.009148407 | **Fanca** | -1.512822036 |
| H2-Ab1 | -1.012510267 | **Padi4** | -1.559250418 |
| Spi1 | -1.01279437 | **Ube2l6** | -1.575842417 |
| Lair1 | -1.022046872 | **Trim10** | -1.725334311 |
| Cd48 | -1.022938848 | **Nr4a3** | -1.744915803 |
| Ccl6 | -1.033292395 | **Alas2** | -1.838940463 |
| Clec4a2 | -1.034299613 | **Ifi206** | -1.927011795 |
| Isg20 | -1.050864189 | **Alcam** | -1.939141258 |
| Slamf9 | -1.056896509 | **Hba-a1** | -2.016402048 |
| Cd74 | -1.065118283 | **Slpi** | -2.190882909 |
| Cd68 | -1.068985937 | **Cxcr5** | -2.215039102 |
| Mgl2 | -1.070726597 | **Slc4a1** | -2.374779065 |
| Bpgm | -1.074345404 | **Hba-a2** | -2.566149469 |
| Slc11a1 | -1.074509333 | **Cxcr2** | -2.889608531 |
| Slamf9 | -1.056896509 | **Tspo2** | -2.982078678 |
| Pld4 | -1.077841498 | **Ifi27l2b** | -3.505870927 |
| Pstpip1 | -1.088503174 | **Alox15** | -3.713626773 |
| Mmp9 | -1.096067239 | **Pvrig** | -4.133852948 |
| Clec4a1 | -1.097355767 | **Tnfsf18** | -4.781439027 |
| Racgap1 | -1.09911279 | **C4a** | -4.789294712 |
| Ccdc92 | -1.099450893 | **Marco** | -5.048140046 |
| Ccl11 | -1.101724435 | **Rab44** | -5.214935477 |
| Serpina3g | -1.103081683 |  |  |

**Immune Genes**

| **Female**  Continued….. | | **Male** |
| --- | --- | --- |
| **Downregulated Gene** | | **Downregulated Gene** |
| **Ssc5d** | -1.155544153 |  |
| **Pilrb1** | -1.162869246 |  |
| **Hba-a1** | -1.173558027 |  |
| **Tnfrsf13b** | -1.179923062 |  |
| **Ccl8** | -1.182650567 |  |
| **Ncf1** | -1.19483589 |  |
| **Clec10a** | -1.215553505 |  |
| **Ly86** | -1.216740179 |  |
| **Pik3ap1** | -1.218579976 |  |
| **Hspa1b** | -1.231541662 |  |
| **Csf2rb** | -1.236723255 |  |
| **Laptm5** | -1.24465392 |  |
| **Adgre1** | -1.249594015 |  |
| **Ube2l6** | -1.265948788 |  |
| **Ciita** | -1.275776301 |  |
| **Fcna** | -1.359178142 |  |
| **Cd300lb** | -1.361360356 |  |
| **Cd19** | -1.3957274 |  |
| **Epb42** | -1.40365059 |  |
| **Smpdl3b** | -1.442841061 |  |
| **Wfdc17** | -1.467612205 |  |
| **Ccr1** | -1.493310161 |  |
| **Fcgr4** | -1.507737631 |  |
| **Hba-a2** | -1.529287757 |  |
| **Alas2** | -1.565451483 |  |
| **Cebpb** | -1.613891546 |  |
| **Il18rap** | -1.815030253 |  |
| **Igkv4-91** | -1.843472972 |  |
| **Oas3** | -1.851808103 |  |
| **Milr1** | -1.936537749 |  |
| **Nectin4** | -2.039084784 |  |
| **Aurkb** | -2.049472392 |  |
| **Clec5a** | -2.073962643 |  |
| **Igkv8-24** | -2.926358517 |  |
| **Ncr1** | -3.552328977 |  |
| **Ighv8-8** | -4.249125228 |  |
| **Ighv9-3** | -4.494770799 |  |
| **Ighv9-1** | -4.995205339 |  |
| **Igkv3-7** | -6.183017903 |  |
| **Batf2** | -1.109570189 |  |

**Cytoskeleton Genes**

| **Female** | | **Male** | |
| --- | --- | --- | --- |
| **Gene Symbol** | **Log2FC** | **Gene Symbol** | **Log2FC** |
| **Upregulated Gene** | | **Upregulated Gene** | |
| **Snap25** | 4.202628595 | **Kif6** | 1.516440195 |
| **Myh8** | 4.120464972 | **Myh7** | 1.402908798 |
| **Rsph10b** | 3.112476172 | **Amph** | 1.163136868 |
| **Tekt2** | 2.87397185 | **Downregulated Gene** | |
| **Hydin** | 2.782751492 | **Ccdc15** | -1.086462243 |
| **Erich3** | 2.583650905 | **Krt8** | -1.099702399 |
| **Sptbn5** | 2.178926143 | **Sntb1** | -1.190562272 |
| **Lmntd1** | 1.996834502 | **Hspa1a** | -1.226780923 |
| **Alox8** | 1.950791021 | **Abcc3** | -1.267002066 |
| **Neb** | 1.707070934 | **Sass6** | -1.318697819 |
| **Cfap74** | 1.698037516 | **S100a9** | -1.34944444 |
| **Enkur** | 1.490204807 | **Procr** | -1.440977188 |
| **Trim67** | 1.402659538 | **Snca** | -1.488085739 |
| **Dcdc2b** | 1.359419582 | **Akap5** | -1.550047804 |
| **Rp1** | 1.283438581 | **Cmah** | -1.752960772 |
| **Plekha7** | 1.277717318 | **Myoz1** | -2.151414696 |
| **Ttn** | 1.272920799 | **Slc4a1** | -2.374779065 |
| **Dnah8** | 1.264941561 | **Gnat3** | -2.419668981 |
| **Ush1c** | 1.107797584 | **Dnhd1** | -2.868169713 |
| **Pclo** | 1.060884748 | **Cxcr2** | -2.889608531 |
| **Myo5c** | 1.045320279 | **Ak8** | -4.31866328 |
| **Ddx11** | 1.015388085 | **Ttll10** | -4.944113015 |
| **Myh7b** | 1.013502547 | **Cfap58** | -5.091259464 |
| **Downregulated Gene** | | **Cfap44** | -5.091259464 |
| Cd86 | -1.00015766 | **Krt23** | -5.799120097 |
| Cys1 | -1.007318409 |  |  |
| Myo1g | -1.009148407 |  |  |
| Pstpip1 | -1.088503174 |  |  |
| **Racgap1** | -1.09911279 |  |  |
| **Ccdc92** | -1.099450893 |  |  |
| **Hspa1b** | -1.231541662 |  |  |
| **Kcnab2** | -1.237394554 |  |  |
| **Epb42** | -1.40365059 |  |  |
| **Kif2c** | -1.870327002 |  |  |
| **Aurkb** | -2.049472392 |  |  |
| **Tnnc2** | -2.328357832 |  |  |
| **Dnajb13** | -3.205096932 |  |  |
| **Axdnd1** | -4.744907947 |  |  |

**Lipid Metabolisms Genes**

| **Female** | | **Male** | | |
| --- | --- | --- | --- | --- |
| **Gene Symbol** | **Log2FC** | **Gene Symbol** | | **Log2FC** |
| **Upregulated Gene** | | **Upregulated Gene** | | |
| **Cyp2c23** | 3.675912947 | **Apoa1** | 5.867932686 | |
| **Bmpr1b** | 2.640685411 | **Elovl4** | 3.047406665 | |
| **Fut2** | 2.097245167 | **Downregulated Gene** | | |
| **Nudt8** | 2.004477616 | **B4galnt1** | -1.12246194 | |
| **Alox8** | 1.950791021 | **Ltc4s** | -1.125582663 | |
| **Dkk3** | 1.920499315 | **Gpx1** | -1.17654102 | |
| **Dpm1** | 1.435202295 | **Snca** | -1.488085739 | |
| **Bmp2** | 1.250570795 | **Rdh12** | -1.987602359 | |
| **Asxl3** | 1.206137957 | **Elovl6** | -2.398643771 | |
| **Ces1e** | 1.180346317 | **Alox15** | -3.713626773 | |
| **Erbb4** | 1.024349659 | **Mgst2** | -4.106996288 | |
| **Downregulated Gene** | | **Sult1e1** | -5.746642311 | |
| **Cd74** | -1.065118283 | **Acoxl** | -6.382398505 | |
| **Pld4** | -1.077841498 |  |  | |
| **Ltc4s** | -1.113184049 |  |  | |
| **Ugt1a7c** | -1.325618636 |  |  | |
| **Smpdl3b** | -1.442841061 |  |  | |
| **Cyp4f18** | -1.595644025 |  |  | |
| **Degs2** | -2.608518484 |  |  | |

| **Female** | | **Male** | |
| --- | --- | --- | --- |
| **Gene Symbol** | **Log2FC** | **Gene Symbol** | **Log2FC** |
| **Upregulated Gene** | | **Downregulated Gene** | |
| **Sphkap** | 3.13703915 | Cbr2 | -1.153876147 |
| **Fam124b** | 2.130450861 | Gpx1 | -1.17654102 |
| **Nudt8** | 2.004477616 | Hspa1a | -1.226780923 |
| **Erbb4** | 1.024349659 | Hsp90aa1 | -1.483301564 |
| **Downregulated Gene** | | Snca | -1.488085739 |
| Azin2 | -1.215795228 | Alas2 | -1.838940463 |
| Hspa1b | -1.231541662 | Tspo2 | -2.982078678 |
| Mthfd1l | -1.36861859 | Ifi27l2b | -3.505870927 |
| Hk3 | -1.44982065 | Aifm3 | -4.501576765 |
| Alas2 | -1.565451483 |  |  |
| mt-Nd6 | -1.606067956 |  |  |
| Acat3 | -2.088444451 |  |  |

**Mitochondrial Genes**
