## Supplementary Figure 1-3 and Table 1-2 for "Sex-Specific Characterization of a Novel Osteoarthritis-Induced Heart Failure Model in Mice": Table 2.docx

**Table 2: Primary Antibody Dilution Used in Immunoblotting**

| **No.** | **Primary Antibody** | **Dilution** | **Catalogue No.** |
| --- | --- | --- | --- |
| 1 | **ANP** | 1:1000 | GeneTex GTX109255 |
| 2 | **BNP** | 1:1000 | Abcam ab236101 |
| 3 | **Col1/Col3** | Col1- 1:800  Col3-1:800 | Col1-Novus Biologicals NBP1-30054  Col3- Novus Biologicals NB600-594 |
| 4 | **p38** | Phospho p38- 1:500  Total p38- 1:1000 | Pho-Cell signaling tech 9211S  Total- Cell signaling tech 9212S |
| 5 | **p65** | Phospho p65- 1:500  Total p65- 1:1000 | Pho- Cell signaling tech 3031S  Total- Cell signaling tech 8242 |
| 6 | **Lgals4** | 1:1000 | GeneTex GTX127351 |
| 7 | **Ero1** | 1:500 | Abcam AB177156 |
| 8 | **Prdx4** | 1:500 | Abcam ab184167 |
| 9 | **PDI** | 1:500 | Cell signaling tech 3501T |
| 10 | **4-HNE** | 1:500 | Abcam ab46545 |
| 11 | **BiP** | 1:1500 | Abcam ab21685 |
| 12 | **Sirt1** | 1:750 | Abcam ab110304 |
| 13 | **Sirt2** | 1:750 | Cell signaling tech 12650T |
| 14 | **Sirt3** | 1:750 | Cell signaling tech 5490T |
| 15 | **pAMPK** | Phospho AMPK- 1:750  Total AMPK- 1:750 | Pho-Cell signaling tech 2531S  Total-Cell signaling tech 2793T |
| 16 | **Hif1α** | 1:750 | Cell signaling tech 14179T |
| 17 | **Casp3** | 1:750 | Cell signaling tech 9661S |
| 18 | **LC3B** | 1:750 | Abcam [ab192890](https://www.abcam.com/en-us/products/primary-antibodies/lc3b-antibody-epr18709-autophagosome-marker-ab192890) |
| 19 | pATG16L1 / total ATG16L1 | Pho ATG16L1- 1:750  Total ATG16L1- 1:750 | Pho- Abcam [ab195242](https://www.abcam.com/en-us/products/primary-antibodies/atg16l1-phospho-s278-antibody-epr19016-ab195242)  Total- Abcam [ab187671](https://www.abcam.com/en-us/products/primary-antibodies/atg16l1-antibody-epr15638-n-terminal-ab187671) |
| 20 | SQSTM1 | 1:750 | Abcam [ab109012](https://www.abcam.com/en-us/products/primary-antibodies/sqstm1-p62-antibody-epr4844-autophagosome-marker-ab109012) |
| 21 | M6PR | 1:1000 | Abcam [ab124767](https://www.abcam.com/en-us/products/primary-antibodies/m6pr-cation-independent-antibody-epr6599-lysosome-membrane-marker-ab124767) |
| 22 | Ubiquitin | 1:1000 | Abcam [ab134953](https://www.abcam.com/en-us/products/primary-antibodies/ubiquitin-antibody-epr8830-ab134953) |
